# Rhythm Receptive Fields in Striatum of Mice Executing Complex Continuous Movement Sequences

**DOI:** 10.1101/2023.09.23.559115

**Authors:** Kojiro Hirokane, Toru Nakamura, Takuma Terashita, Yasuo Kubota, Dan Hu, Takeshi Yagi, Ann M. Graybiel, Takashi Kitsukawa

**Affiliations:** Graduate School of Life Sciences, Ritsumeikan University, Kusatsu, Shiga, Japan; Graduate School of Frontier Biosciences, Osaka University, Suita, Osaka, Japan; McGovern Institute for Brain Research and Dept. of Brain and Cognitive Sciences, Massachusetts Institute of Technology, Cambridge, Massachusetts, United States

**Keywords:** mouse, basal ganglia, striatum, rhythm, electrophysiology, stepping, body coordination, motor chunking, receptive field, locomotion

## Abstract

By the use of a novel experimental system, the step-wheel, we investigated the neural underpinnings of complex and continuous movements. We recorded neural activities from the dorsolateral striatum and found neurons sensitive to movement rhythm parameters. These neurons responded to specific combinations of interval, phase, and repetition of movement, effectively forming what we term “rhythm receptive fields.” Some neurons even responsive to the combination of movement phases of multiple body parts. In parallel, cortical recordings in sensorimotor areas highlighted a paucity of neurons responsive to multiple parameter combinations, relative to those in the striatum. These findings have implications for comprehending motor coordination deficits seen in brain disorders including Parkinson’s disease. Movement encoding by rhythm receptive fields should streamline the brain’s capacity to encode temporal patterns, help to resolve the degrees of freedom problem. Such rhythm fields hint at the neural mechanisms governing effective motor control and processing of rhythmic information.

## Introduction

Many of our actions, such as playing musical instruments and sports, involve continuous coordinated movements among multiple body parts, requiring the ability to coordinate with precision the timing and position of these various body parts. However, selecting an optimal state from an enormous number of possible combinations of joint angles, muscle contraction states and other parameters makes it the extremely difficult problem known as the “combinatorial explosion”^1^. This problem arises due to increased degrees of freedom in movement determination and becomes worse for continuous movements, because planning movements over longer periods of time using optimization strategies becomes exponentially more difficult with increases in time^2, 3^. Therefore, it should further be difficult to determine the position of body parts in continuous movement sequences. Detailed mechanisms for the execution of such movements remain unclear.

In most situations, continuous movements are executed as repetitive movements. One possible way to reduce the degrees of freedom in continuous movements should be to fix the cycle of the repetitive movement, as demonstrated in equiperiodic tapping task^4^. Another strategy involves establishing fixed coordination patterns among body parts, demonstrated in bimanual coordination tasks^5^ in which the phase relationship between the periodic movements of the two arms became stable with learning. These findings suggests that rhythm, interval and phase could control the coordination of the simple continuous movements that can be executed by repeating movements with a fixed interval. For more complex movements, which require constantly changing repetition cycle and positional relationship of body parts, we found that specific movement runs spanning several footsteps emerged as mice were trained on complex running patterns required for rewarded performance in the step-wheel task^6–11^. In these runs, the timing of footsteps became stable from trial to trial. We denoted these stable regions as “chunks”^8^, in reference to classic work in the motor system^12, 13^. These chunks were, in particular, characterized by the stability of both interval and phase, which reflect the rhythm of multiple body parts. The discovery that chunks observed within complex movements were established by rhythm supports the idea that rhythm contributes to reducing computational complexity. Based on these findings, we concluded that the coordination of complex and continuous movements could be governed by rhythm parameters. This is intuitive from our daily experience, either as producers of rhythms as in singing or dancing, or as viewers of rhythms such as present in many sports, but here yielded to quantitative analysis. These abilities suggest the presence of neural mechanisms for perceiving and processing rhythms in complex and continuous movements.

However, the brain mechanisms involved in executing complicated rhythmic movements remain elusive. To address this issue, we used the step-wheel^8, 9^ to record neural activity in the striatum in mice running on differing peg-patterns and searched for neurons with activity related to rhythm parameters. We considered that if coordination of sequential movements is controlled by rhythm parameters, then there might exist neurons that represent intervals, phases, and repetitions. In an earlier study, by early gene expression patterns, we identified the activation of neurons in the dorsolateral striatum of mice after running in the step-wheel^7^, consonant with many lines of evidence supporting the involvement of the striatum in the control of movements including locomotion^11, 14–19^. Patients with Parkinson’s disease (PD) have problems with voluntary movements leading to bradykinesia (slowness of movement) and ultimately akinesia (difficulty in initiating movement) due to striatal dysfunction^20–24^and a hallmark is that motor rhythm is lost in patients suffering from PD^25, 26^ and that patients are aided by external rhythm experience. For these reasons, we aimed to determine the precise firing patterns of the striatal neuron that develop during the acquisition of complex continuous stepping task.

We recorded neural activity from the dorsolateral striatum of mice running in the step-wheel and analyzed the activity patterns of neurons, mainly the medium spiny neurons (MSNs) that constitute as much as 90% of all striatal neurons. We found striatal MSNs that were responsive to peg touching, water drinking, and approach/leave of the water spout. Many peg-touch-responsive neurons exhibited responses as well to specific stepping intervals, phases of stepping, or to numbers of repetition, most of which had combinatorial responses to all three of these rhythm parameters. By contrast, parallel sensorimotor neocortical recordings identified little such combinatorial responsivity. These findings suggest that movements may be encoded in the striatum according to rhythm parameters including interval, phase and repeat patterns in a combinatory form that we here designate as *rhythm receptive fields*.

## Results

The Ferris-wheel like step-wheel was used to train mice to perform complex running movements. Mice learned to step on pegs for their footholds as the wheel was rotated. They were first exposed to a regular, simple peg-pattern to allow them to habituate to the wheel, and then were given a complex peg-pattern to train them to step with various cycles and limb coordination constraints (Fig. 1A–C). Mice were trained to run at a constant speed in the vicinity of a water spout to drink water as a reward (detected by the infrared sensor), matching their running speed to the wheel’s rotation speed, which was kept constant by computer control. The mice were water restricted, and were given a continuous and constant reward throughout the running period. We recorded the touch timing of forelimbs to pegs and the activity of 646 units in 15 mice running in the step-wheel, focusing on the dorsolateral part of striatum (Fig. 1D). Fig. 1E shows the neural activity recorded from one such unit in a running mouse, aligned to one of the turn markers in one rotation of the wheel (two trials). All recordings were performed after at least three weeks of training on each peg-pattern, which gave mice sufficient time to finish learning the peg-pattern and acquire stable running in the wheel.

**Figure 1.**
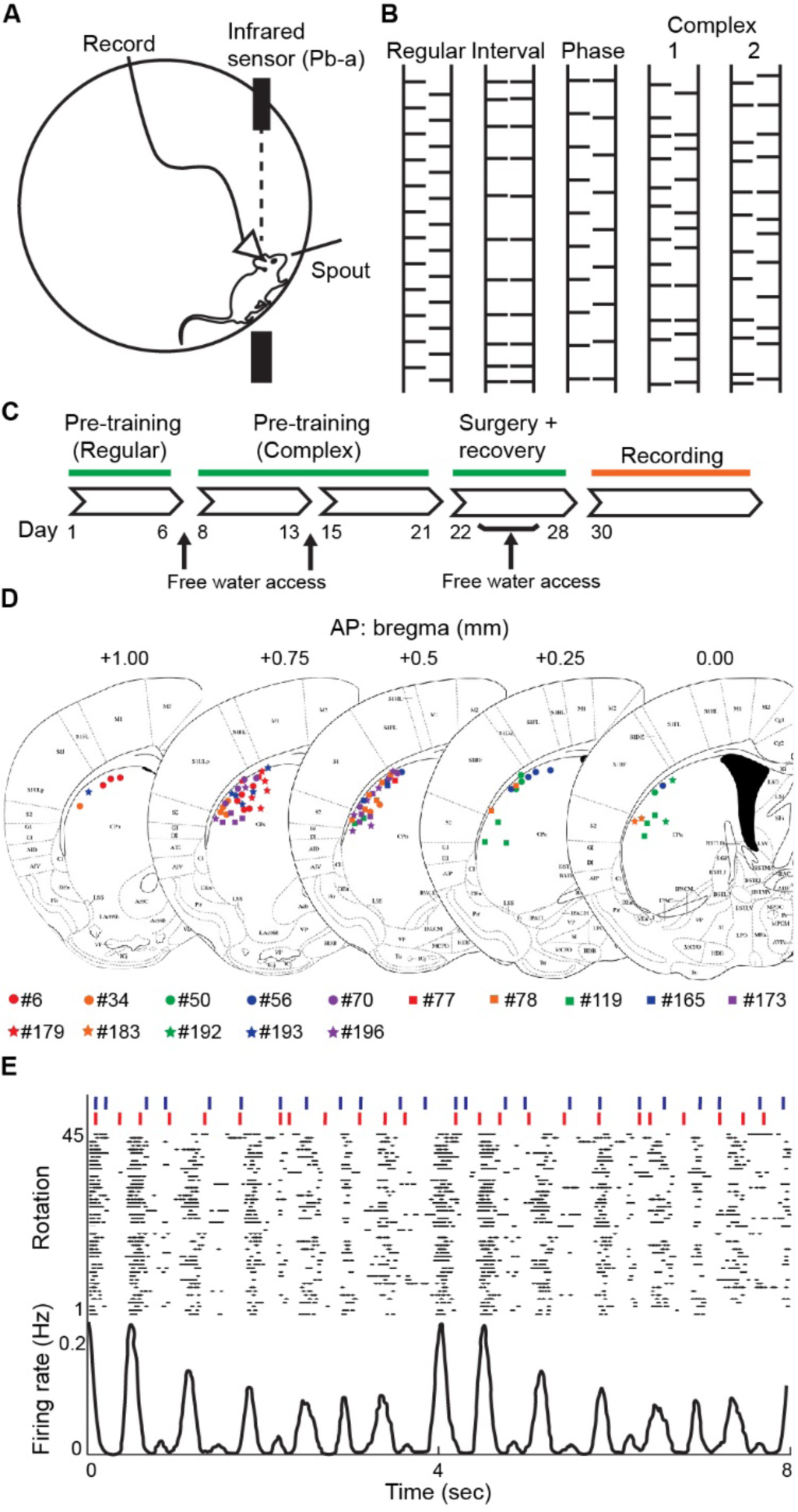
Striatal neurons showed a phasic activity during running in the step-wheel. A. The step-wheel, shown in a schematic diagram. Pegs that serve as footholds for mice were placed on the circumferential surface of the step-wheel. The wheel is motor-driven and rotates at a constant speed. During running and recording experiments, mice were equipped with a head stage containing eight tetrodes for neural recordings. An infrared sensor was positioned to detect the mice approaching to and leaving from the water spout. The voltage sensor on the water spout detects mouth movements during licking, while touch voltage sensors connected to all pegs detect the timing of left and right limb touches to the pegs. B. Peg-patterns used in this study. Four types of patterns were used: a regular pattern of alternating left and right pegs (Regular), a pattern in which only the interval between pegs changes gradually (Interval), a pattern in which only the positional relationship between the left and right pegs changes gradually (Phase), and an irregular pattern (Complex). There were four varieties of complex peg-pattern used in this study, but two of them were illustrated. C. Timeline of the pre-training session, surgery, recovery session, and recording session. D. Final locations of tetrode tips for recording sites. Colors indicate recording sites from different mice. E. Representative single-unit activity recorded from the dorsolateral part of the mouse striatum while running on the complex peg-pattern. Peg-pattern (top), raster plot of firing events in a striatal neuron (middle) and firing rate of a striatal neuron (bottom) are illustrated.

To identify the response characteristics of neurons in detail, we analyzed the correlation of the spike activity to the timing of left and right peg touches, licking recorded by voltage sensors, and approaches to or leaving from the water spout as detected by an infrared sensor. All neural activities recorded were used in the analyses. We found neurons showing phasic activity before and after the limb touches to pegs (Fig. 2A), and neurons showing phasic activity when aligned by the timing of licking (Fig. 2B). Furthermore, by aligning to the timing of approaching to the spout (blockade of the infrared sensor) or to leaving the spout (recovery of the infrared sensor), we found neurons that increased firing rates before mice approached to the spout (Fig. 2C, blue) or after the mouse left the spout (Fig. 2C, red). These findings indicate that the dorsolateral striatum contains neurons responsive to peg touches, licking, and approaching to or leaving from the water spout.

**Figure 2.**
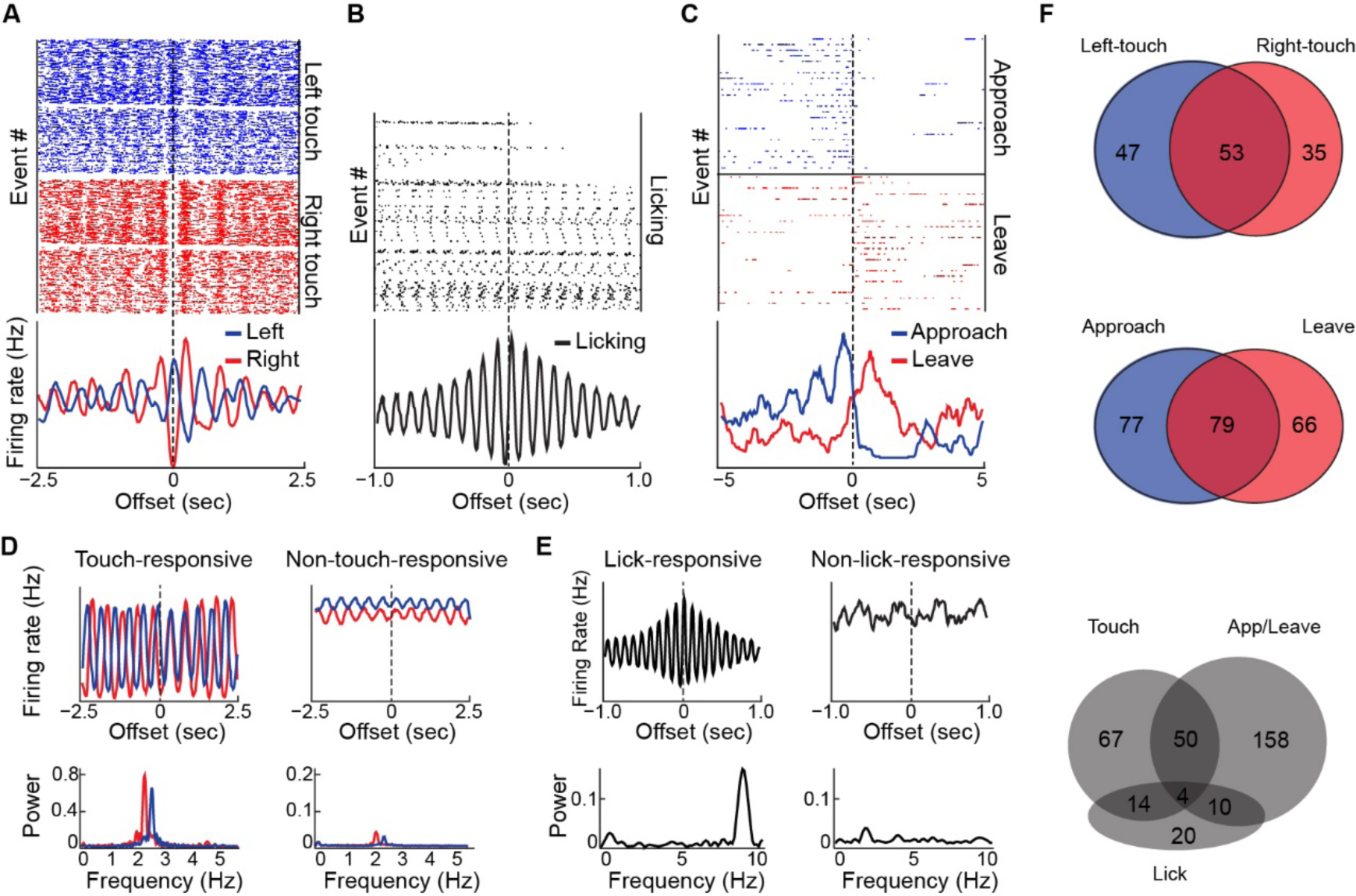
Striatal MSNs responded to each movement. A. Striatal neural activity in response to peg touches. Raster displays (top) and cross-correlograms (bottom) of neural activity aligned with respect to the timing of left (blue) and right (red) forelimb touch to the peg. Cross-correlograms were made with the bin size of 10 msec and smoothing window of 100 msec. B. Striatal neural activity in response to licking. Raster display (top) and cross-correlogram (bottom) of neural activity aligned to the time when the mouse put their mouth on the water spout. Cross-correlograms were made with the bin size of 4 msec and smoothing window of 20 msec. C. Striatal neural activity in response to approaching and leaving the water spout. Raster displays (top) and cross-correlograms (bottom) of neural activity aligned to the time of approach to (blue) and leaving from (red) the water spout. Cross-correlograms were made with the bin size of 20 msec and smoothing window of 200 msec. D and E. Cross-correlograms aligned to the peg touch (D, top) and licking (E, top), and spectrograms calculated from the cross-correlogram for touch timing (D, bottom) and lick timing (E, bottom). Representative touch-responsive MSN (D, left), non-touch-responsive MSN (D, right), lick-responsive MSN (E, left), and non-lick-responsive MSN (right) are shown. For the calculation of the spectrogram, we used original cross-correlogram data, which was not smoothed over. F. The number of MSNs responding to left and right peg touches (top), to approach to and leaving from the spout (middle), and to touch, lick, and approach to/leave from the water spout (bottom).

To identify neurons that responded to touches or licking, we performed Fourier analyses. If a neuron responded to touches or licking, the firing rate should match to the frequency of the event. The cross-correlograms of neural activities aligned by touches (Fig. 2A) or licking (Fig. 2B) were Fourier transformed, and the frequency power was calculated (Fig. 2D, E). We classified neurons according to the power of frequency band corresponding to touches or licking using non-regular peg-patterns (2–3.5 Hz and 7–11 Hz, respectively; see Fig. S1 and STAR Methods for detail).

We identified 100 left-touch-responsive and 88 right-touch-responsive neurons, in which 53 neurons responded to both left and right touches (Fig. 2F, top). The number of neurons that responded to left touches was higher than those responded to right touches, probably because they were recorded from the right striatum. We identified 48 lick-responsive neurons. Touch- and lick-responses were shared by 18 neurons, which corresponded to 13.3 % of touch-responsive neurons and 29.2 % of lick-responsive neurons (Fig. 2F, bottom). The actual overlap might even be less, as licking itself was frequently time-coordinated with touches^8^, increasing the frequency components of the other in each other’s preferred frequencies. We also identified 156 approach-responsive and 145 leave-responsive neurons by comparing firing rates before and after the infrared sensor blockade or release (Fig 2F, middle). Of these, 79 neurons exhibited both approach and leave responses, as exemplified by the neuron shown in Fig. 2C.

Neurons were further classified as MSNs and as fast-spiking interneurons (FSIs) based on recorded waveforms^27–29^. We identified 349 MSNs (Fig. S2A, red) and 91 FSIs (Fig. S2A, blue) out of the total 646 units recorded. The yields of MSNs was not commensurate with what otherwise would be expected values (∼ 90% of total), due to our need to set high thresholds for their identification (See STAR Methods for detail). The striatal neurons identified as touch-responsive, lick-responsive, and approach/leave-responsive were further classified into MSN, FSI, and other categories (Table 1, Fig. S3). We found that the contribution of FSIs was quite low for the touch-responsive neurons compared to lick- and approach/leave-responsive neurons.

**Table 1.**
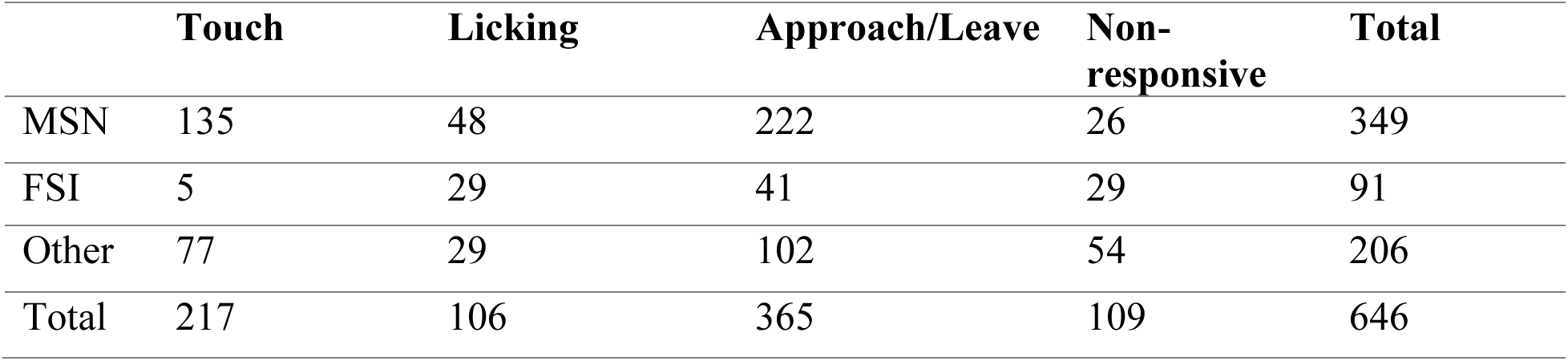
Number of neurons with responsiveness to each movement recorded from all mice used in this study (n = 15 mice).

|  | **Touch** | **Licking** | **Approach/Leave** | **Non-responsive** | **Total** |
| --- | --- | --- | --- | --- | --- |
| MSN | 135 | 48 | 222 | 26 | 349 |
| FSI | 5 | 29 | 41 | 29 | 91 |
| Other | 77 | 29 | 102 | 54 | 206 |
| Total | 217 | 106 | 365 | 109 | 646 |

**Table 2.** Number of neurons with responsiveness to each movement recorded from mice running all the interval-changing, phase-changing and complex peg-patterns on the same day (n = 13 mice).

|  | **Touch** | **Licking** | **Approach/Leave** | **Non-responsive** | **Total** |
| --- | --- | --- | --- | --- | --- |
| MSN | 112 | 42 | 185 | 23 | 301 |
| FSI | 5 | 21 | 36 | 21 | 69 |
| Other | 63 | 32 | 88 | 44 | 177 |
| Total | 180 | 95 | 309 | 88 | 547 |

### Touch-responsive MSNs increased the firing rate at a particular peg touch

We focused on touch-responsive MSNs to investigate what encoding might relate to motor coordination exhibited during the complex continuous stepping. The touch-responsive MSNs were not equally responsive to all peg touches, but only to certain peg touches, which in turn meant that these MSNs did not simply respond to any touch at all. Fig. 3A shows neural activities of two touch-responsive MSNs (units #1 and #2) aligned to the turn marker (start of the trial, half a full cycle of the wheel). Each unit increased its firing rate prominently in two or three distinct locations in the peg-pattern (Fig. 3A). Given that one trial consists of 24 pegs, mice should have touched pegs at least 20 times in a trial.

**Figure 3.**
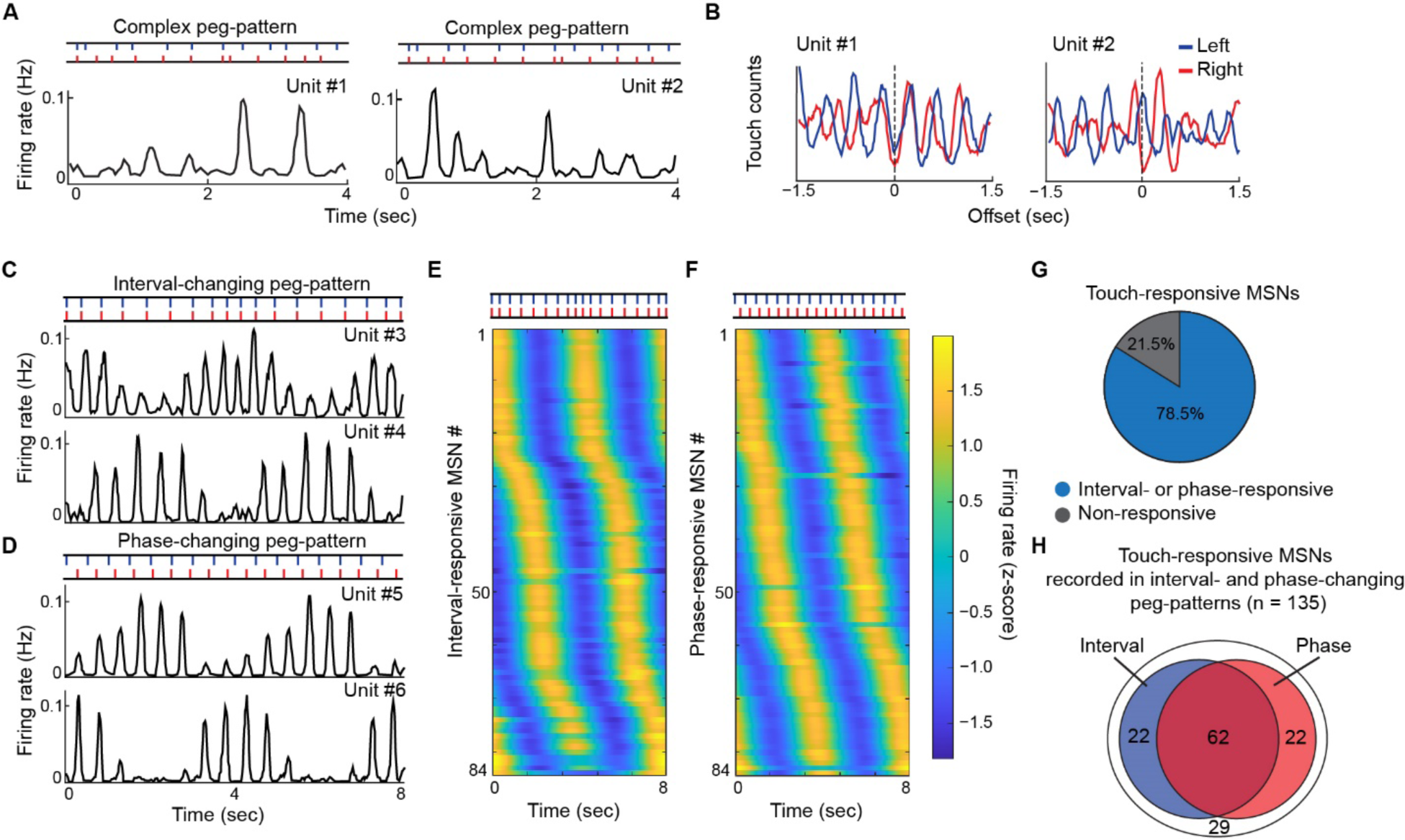
Interval- and phase-responsive neurons were found in the striatum. A. Neural activity of two MSNs (units #1 and #2) during running on the complex peg-pattern. Neural activities were aligned to the turn marker timing. B. Cross-correlogram of peg touches aligned to spike timings of the two MSNs shown in A. C. Peg placement of interval-changing peg-pattern for two trials (top) and firing rates of two MSNs (units #3 and #4) responsive to intervals (middle and bottom). D. Peg placement of phase-changing peg-pattern for two trials (top) and firing rates of two MSNs (units #5 and #6) responsive to the relative position of the left and right limbs (middle and bottom). E and F. Heatmaps visualizing the envelope of z-scored firing rate during running on the interval-changing (E) and phase-changing (F) peg-patterns. Activity of all the interval-responsive (n = 84, E) and all phase-responsive (n = 84, F) MSNs were sorted based on their peak position of firing rate within the pattern. G. Pie chart showing the proportion of touch-responsive MSNs responded to any of interval or phase. H. Venn diagram showing the number of touch-responsive MSNs identified as interval-responsive (blue) and phase-responsive (red) MSNs.

To examine the relationship between touch and MSN firing in more detail, the touch times of the left and right limbs were aligned to the spike times. The cross-correlograms of left and right touches aligned to spike times of these units demonstrated differences in responsiveness (Fig. 3B). For example, the relative timing of the left touch to the spikes was different for units #1 and #2. In addition, the relative timings of the left and right touches around the center of the cross-correlogram (peaks of blue and red lines) were also different; the left and right limbs touched almost simultaneously around the spike timing for unit #1, whereas the left and right limbs touched alternately around the spike timing for unit #2. Finally, the intervals between the peaks to the adjacent left touch (blue) were longer for unit #2 than in unit #1. These differences indicate that the interval between peg touches and the timing of the left and right touches (left-right difference) likely produced the differences in firing patterns of these units, as shown in Fig. 3A. This result suggested the possibility that the activity of touch-responsive MSNs might depend on the interval and left-right difference of peg touches.

### Touch-responsive MSNs responsive to the intervals of touch cycle

To address this possibility, we next asked whether the activity of touch-responsive MSNs changed their firing patterns in relation to the width of touch intervals or the relative left-right limb positions by establishing two different peg-patterns for these purposes: interval-changing and phase-changing peg-patterns. In the interval-changing pattern, peg intervals gradually changed, so that mice experienced a variety of peg intervals (Fig. 1B, Interval). In the phase-changing peg-pattern, timing differences between the left and right pegs gradually changed, so that the mice experienced a variety of left-right coordination patterns (Fig. 1B, Phase). To determine whether each MSN could possess multiple responsiveness to touch intervals, left-right limb positions, or others, MSNs that were recorded from mice required to run on these two peg patterns and at least one complex pattern within a single day and that, in addition, showed firing rates of more than 0.5 spikes/sec in all of sessions were used for the analyses.

First, to test the possibility that touch-responsive MSNs responded to the interval of peg touches, we analyzed neural activity recorded when mice were running on the interval-changing peg-pattern. We found MSNs that increased their firing rates at specific locations on the interval-changing peg-pattern (Fig. 3C). The firing patterns of two exemplar units (#3 and #4) are illustrated with the interval-changing peg-pattern: unit #3 increased firing rate at relatively narrow intervals and decreased firing rate at broader intervals (Fig. 3C, top trace). By contrast, unit #4 increased its firing rate at broader intervals and decreased it at narrower intervals (Fig. 3C, bottom trace). For the full data set, we defined as interval-responsive MSNs the touch-responsive MSNs that changed their firing rates depending on the interval between peg touches in the interval-changing peg-pattern. Specifically, a discrete Fourier transform (DFT) was performed on the envelope of peaks in firing rates over the span of two trials, and spectrograms were obtained. The spectrograms were then z-scored, and touch-responsive MSNs that showed a specified z-scored power (> 1.8, see STAR Methods for detail) at 0.25 Hz frequency were identified as the interval-responsive MSNs. Note that the DFT was performed on the envelope of peaks in firing rates over the span of one rotation (8 sec). There are two trials in one rotation, and the interval-responsive MSNs increased the firing rate at one specific location within a trial. Thus, the increase or decrease was repeated twice per wheel rotation (2 trials).

Among the 135 touch-responsive MSNs recorded, 84 interval-responsive MSNs were identified. We sorted the interval-responsive MSNs by the location of the maximal firing rate over the peg-pattern and found that the optimal intervals differed from neuron to neuron. A slightly larger proportion of MSNs were active at the narrowest and widest intervals compared to the locations between the two extremes, but almost all intervals that appeared in the interval-changing peg-pattern were covered by these interval-responsive MSNs (Fig. 3E). Interval-responsive MSNs that preferred wider or narrower intervals than the intervals that appeared in this peg pattern may have been active at either extreme.

### Touch-responsive MSNs responsive to the timing of left and right touch cycle

Next, we analyzed neural activity recorded when mice were running on the phase-changing peg-pattern to examine the possibility that touch-responsive MSNs respond to the left-right difference in limb position. In the phase-changing peg-pattern, the peg intervals were slightly wider on one side (left or right) but identical in each side, thereby generating different left-right relationship for each step while minimizing the effect of the interval. The left-right difference was defined as small when the left and right pegs were aligned face-to-face and defined as large when the left and right pegs were aligned alternately.

We found MSNs with increased firing rates at specific locations in the phase-changing peg-pattern (Fig. 3D). As an example, for unit #5, the firing rate increased when the left-right difference was small and decreased when the left-right difference was large (Fig. 3D, top trace). By contrast, for unit #6, the firing rate increased when the left-right difference was large and decreased when the left-right difference was small (Fig. 3D, bottom trace). The relative position of the left and right limbs can be considered as the touch timing of the contralateral limb within the cycle of the referenced side’s limb. Thus, the relative positions can be expressed as a phase (0–2π), which is the reason why the units for which the firing rates depended on the phase in the phase-changing peg-pattern were defined as phase-responsive MSNs. As used in determining the interval-responsive MSNs, spectrograms were obtained and then z-scored. MSNs that showed a specified z-scored power > 2.0 (see STAR Methods for detail) at 0.25 Hz frequency were identified as the phase-responsive MSNs.

Among the 135 touch-responsive MSNs, we identified 84 phase-responsive MSNs. When these MSNs were sorted by the location of maximal firing rate within the peg-pattern, each MSN showed an optimal responsive phase, and the distribution of optimal responsive phases for these MSNs covered all regions of the phase-changing peg-pattern (Fig. 3F). A majority (78.5%) of touch-responsive MSNs were classified as interval- and/or phase-responsive MSNs (Fig. 3G), and 45.9% of the touch-responsive MSNs responded to both the interval and the phase. In addition, over a half of the interval-responsive MSNs showed phase responsiveness, and similarly, over a half of the phase-responsive MSNs showed interval responsiveness (Fig. 3H). These results suggest that many interval- and phase-responsive MSNs are likely to belong to a single group of neurons responsive to both interval and phase rather than belonging to separate groups of interval- and phase-sensitive neurons. Thus, our findings indicate that most touch-responsive MSNs in the dorsolateral striatum have interval and phase-related responses (Fig. 3G).

### Touch-responsive MSNs fired at a specific phase within the touch cycle

Having observed that the relative timing of touches on the right and left limbs to spikes could differ for two MSNs (Fig. 3B), it seemed likely that the phase-responsive MSNs that were responsive to the positional relationship might have responded to some relationship between left and right limb positions. Possible mechanisms for this relationship could include the responsiveness to absolute or relative time from the preceding touch or to the subsequent touch. We set up six hypotheses (Fig. S4A) and performed a decoding analysis of the neural activity to narrow down the most likely mechanisms. We made six models corresponding to these six hypotheses, using peg touching and neural activity data recorded in sessions run on the complex peg-pattern. We then predicted the firing rate of the same phase-responsive MSNs on the phase-changing peg pattern using the models and compared them to the actual firing rate. The most plausible model was estimated by the accuracy of predictions by introducing peak error (Fig. S4C) and Jensen-Shannon divergence (JSD, Fig. S4D) as error indicators. As a result of comparing the prediction accuracy among the six models, the accuracy of the prediction data using model 1 was the best for both indicators (Fig. 4A: p < 0.05 by one-factor ANOVA; p < 0.05 by Bonferroni test). This result suggested that phase-responsive MSNs have an optimal phase to fire (relative time, not absolute time) in touch cycles of each limb as shown in Fig 4B, which illustrates phase response probabilities of firing in a cycle for each limb (Fig. 4B, bin = 0.1π, where one cycle is 2π).

**Figure 4.**
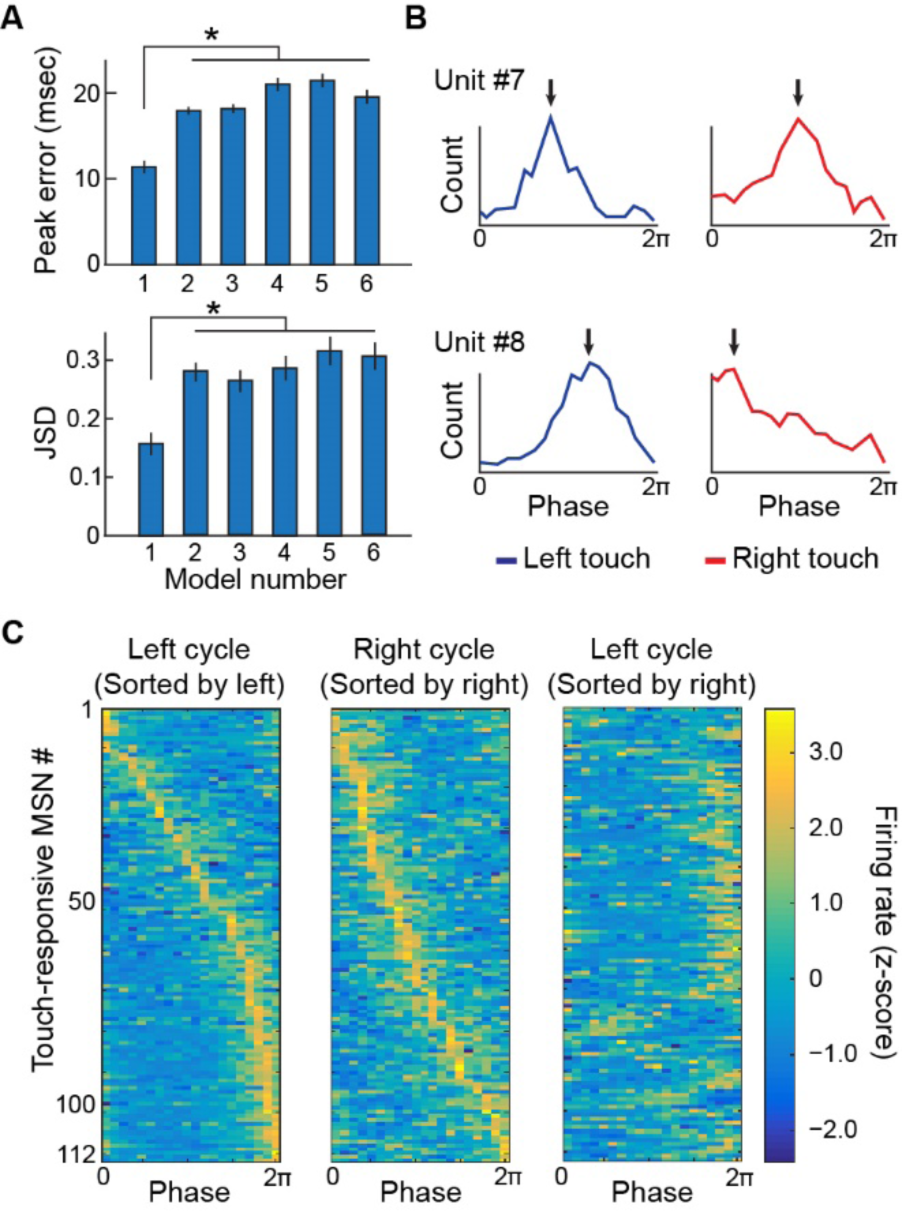
Touch-responsive MSNs responded to a specific phase within a touch cycle. A. Prediction accuracies of six models evaluated by peak error and JSD (n = 112 units, p < 0.05 by one-factor ANOVA; *p < 0.05 by Bonferroni test). See Fig. S4 for the detail of each model. B. Histogram of spike counts that occurred within each of the left (blue) and right (red) touch cycles for two touch-responsive MSNs (units #7 and #8) during running on the complex peg-pattern. Arrows indicate the position of peak firing rate for each MSN within each of the left and right touch cycles. C. Heatmaps of the z-scored firing rates of touch-responsive MSNs for the left and right touch cycles, sorted based on their peak positions within each left (left) and right (middle) touch cycle. The heatmap at right shows z-scored firing rates within the left touch cycle, sorted based on their peak positions within right cycle. These analyses were performed on touch-responsive MSNs recorded from mice that ran both the phase-changing and complex peg-pattern (n = 112 units in 13 mice).

We aligned the optimal phases of each touch-responsive MSN by its peak position in each left and right cycle (Fig. 4C, left and middle). The optimal responses phase of the MSNs covered almost all phases: 0 to 2π. In addition, the optimal response phases of MSNs in the left cycle seemed to be independent of those in the right cycle (Fig. 4C, right), indicating that all positional relationships of left and right limb movement can be represented by the MSNs.

### Touch-responsive MSNs are responsive to the number of repetitions of touch cycle

Considering that the peg touches by the left and right limbs seemed to be approximately equally spaced when aligned by spike timings (Fig. 3A, B), and that most touch-responsive MSNs were interval-responsive MSNs (Fig. 3H), we asked whether MSN activity might be modulated by multiple repetitive touch cycles.

We examined the relationship between the number of touch repetitions and the firing rate of these MSNs recorded from mice running on the complex peg-pattern by putting together five equally spaced touch windows and one spike window as a ‘window set’ and then slid the set through the session time (Fig. 5A). By counting the number of consecutive touch windows in which peg touch events were found, we could quantify the number of repetitions of touch cycle at each time point. The touch windows were equally spaced in the time direction, which made it possible to quantify the periodicity of the movement by counting the number of consecutive windows in which the touch events occurred (repetitive touch cycle). The firing rates at each time point were calculated by counting the number of spikes that occurred in the spike window. The intervals between adjacent touch windows in a window set were moved in 10-msec step intervals from 300 to 500 msec (Fig. 5B), and the positional relationship between the spike windows and the touch windows was set at phases from −π to π in 0.2π steps (Fig. 5B). With the 3^rd^ touch window as the reference point (0π, center), one spike window was fixed in the period between −π to π around the reference point. This analysis was performed on all touch-responsive MSNs recorded in the striatum.

**Figure 5.**
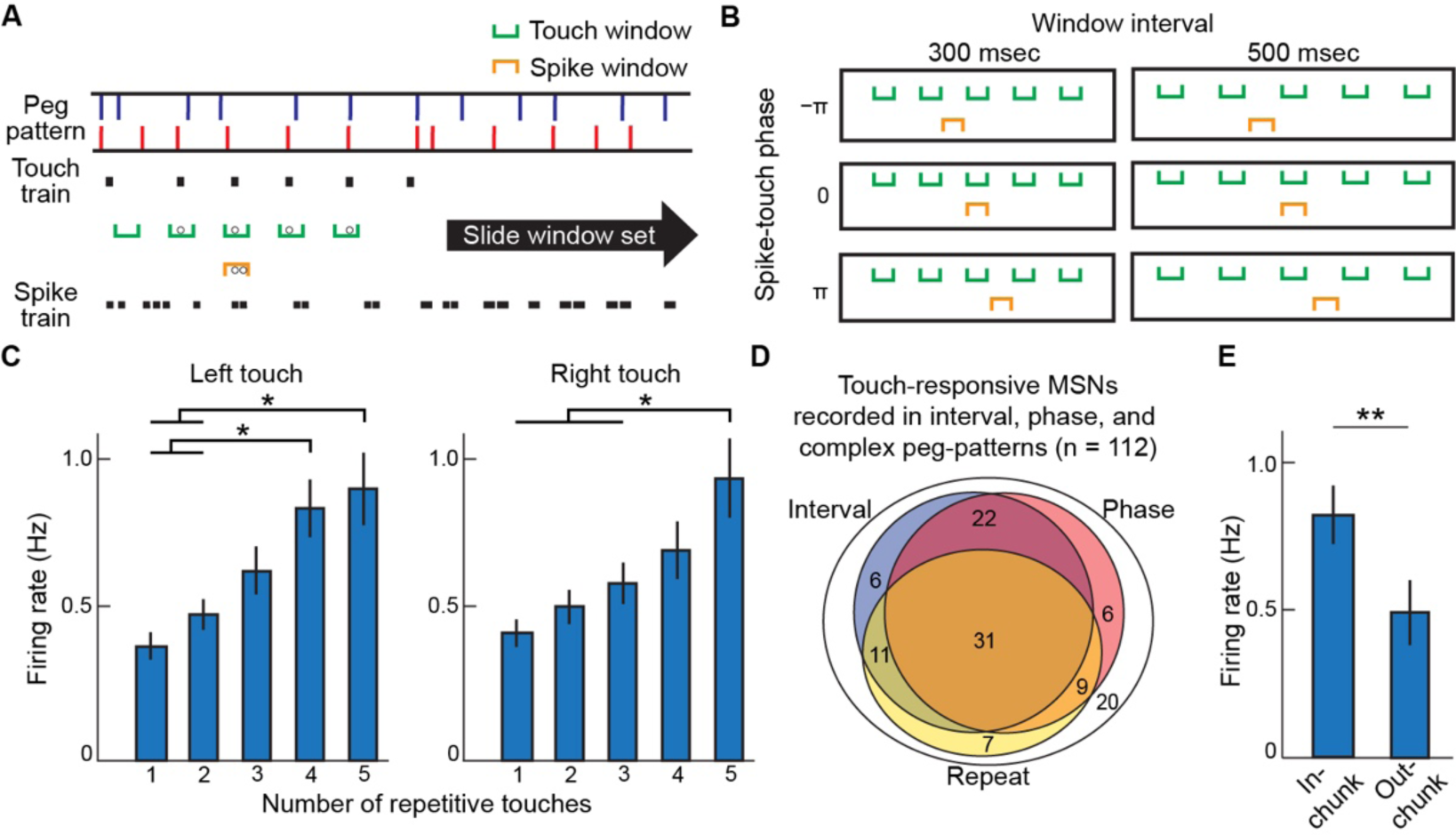
Touch-responsive MSNs increased the firing rate with repetition. A. Schematic diagram illustrating window analysis. A window set contained five evenly spaced touch windows for the detection of touches at a regular interval and a spike window for the detection of firing events. B. Evenly spaced touch windows were set at various intervals (300–500 msec intervals, every 10 msec), with spike windows in various phases before and after the third touch window (−π to π, every 0.2π). C. Average firing rate of all touch-responsive MSNs, recorded from mice that ran on the phase-changing, interval-changing and complex peg-pattern, in relation to the number of repetitive touch cycles (n = 112 units, p < 0.05 by one-factor ANOVA; *p < 0.05 by Bonferroni test). D. The number of interval-, phase- and repeat-responsive MSNs among the touch-responsive MSNs recorded from mice that ran on the phase-changing, interval-changing and complex peg patterns on the same day and used for window analysis. E. The firing rate of the repeat-responsive MSNs calculated separately for inside and outside the chunk (**p = 4.6×10^-6^ by two-tailed t-test).

The average firing rate increased as the number of repetitive touch cycles increased (Fig. 5C, p < 0.05 by one-factor ANOVA; p < 0.05 by Bonferroni test). This result indicates that touch-responsive MSNs changed their firing rates based on the repetition number of the touches in addition to the interval or phase of touches. Such repeat-responsive MSNs were defined by performing linear regression to estimate the relationship between the number of repetitions and the firing rate for each MSN. Regression coefficients (*β*) and coefficients of determination (Rsq) obtained by linear regression were used as the criteria for the identification of the repeat-responsive MSNs. The number of repeat-responsive MSNs are shown in Fig. 5D together with the interval- and phase-responsive MSNs. Among the interval- and/or phase-responsive MSNs, 60.0% (51/85 units) responded to the repeat, indicating that many interval- and phase-responsive MSNs responded to the repetition of touch cycles. Similarly, almost all repeat-responsive MSNs (51/58 units, 87.9%) responded to the interval and/or phase. This finding indicates that most interval-, phase- and repeat-responsive MSNs are likely to belong to a single group of neurons with overlapping responsiveness to these parameters.

Repeat-responsive MSNs increased their firing rates when the movement was repeated in the same interval, that is, according to a rhythm. Our previous study demonstrated that the rhythm of each body part is stable within behaviors that are acquired as motor chunks^8^. We asked whether the interval-, phase- and repeat-responsive MSNs also responded according to this motor chunking. We identified motor chunk regions using the chunk identification method used in Hirokane et al.^8^ (see STAR Methods), and we compared the firing rates of interval-, phase- and repeat-responsive MSNs inside and outside of the chunk. We found that the firing rate of these MSNs was higher inside of the chunks (Fig. 5E, p = 4.6 × 10^-6^ by two-tailed t-test). This result suggests that MSNs that responsive to interval, phase, and repetition could be involved in the execution of rhythm chunks by increasing their firing rates inside the chunked regions acquired through experience.

### Repeat-responsive MSNs increased their firing rates with repetition of specific interval and phase of movements

Most touch-responsive MSNs exhibited responsiveness related to interval, phase, or repetition (Fig. 5D). This finding suggests that the touch-responsive MSNs have their optimal intervals and phases, respectively. However, since the peg-pattern used for the identification of interval-responsive MSNs (i.e., interval-changing peg-pattern) had a fixed phase (0 pi) and the peg-pattern used for the phase-responsive MSNs (i.e., phase-changing peg-pattern) had fixed intervals for left and right limbs, it is not obvious whether the MSNs have optimal interval and phase at the same time. In addition, it remained possible that the MSNs might have been active outside of their optimal interval and phase so long as the touches were repeated. To address these issues, we next examined the responsiveness to every interval and phase combination under the high repetition state for individual MSNs recorded from mice running on the complex peg-pattern. We defined a high repetition state to be the condition in which touches were repeated for four cycles or more, and the low repetition state to be the condition in which touches were repeated for fewer than four cycles at a given interval and phase. The repeat responsiveness of each MSN was analyzed by a series of window sets with different interval and phase pairs. The interval-phase firing characteristics of two representative repeat-responsive MSNs (units #9 and #10) under the high repetition state are depicted in heatmaps with interval and phase as, respectively, the vertical and horizontal axes (Fig. 6A). We found that the repeat-responsive MSNs responded to the repetitions of optimal intervals and phases that differed from neuron to neuron (Fig. 6B). For many MSNs, the interval-phase firing characteristics significantly differed between the high repetition state (Fig. 6B) and low repetition state (Fig. S5A). The firing rates under the low repetition state were low as a whole and exhibited fewer activity preferences for interval and phase (Fig. S5A).

**Figure 6.**
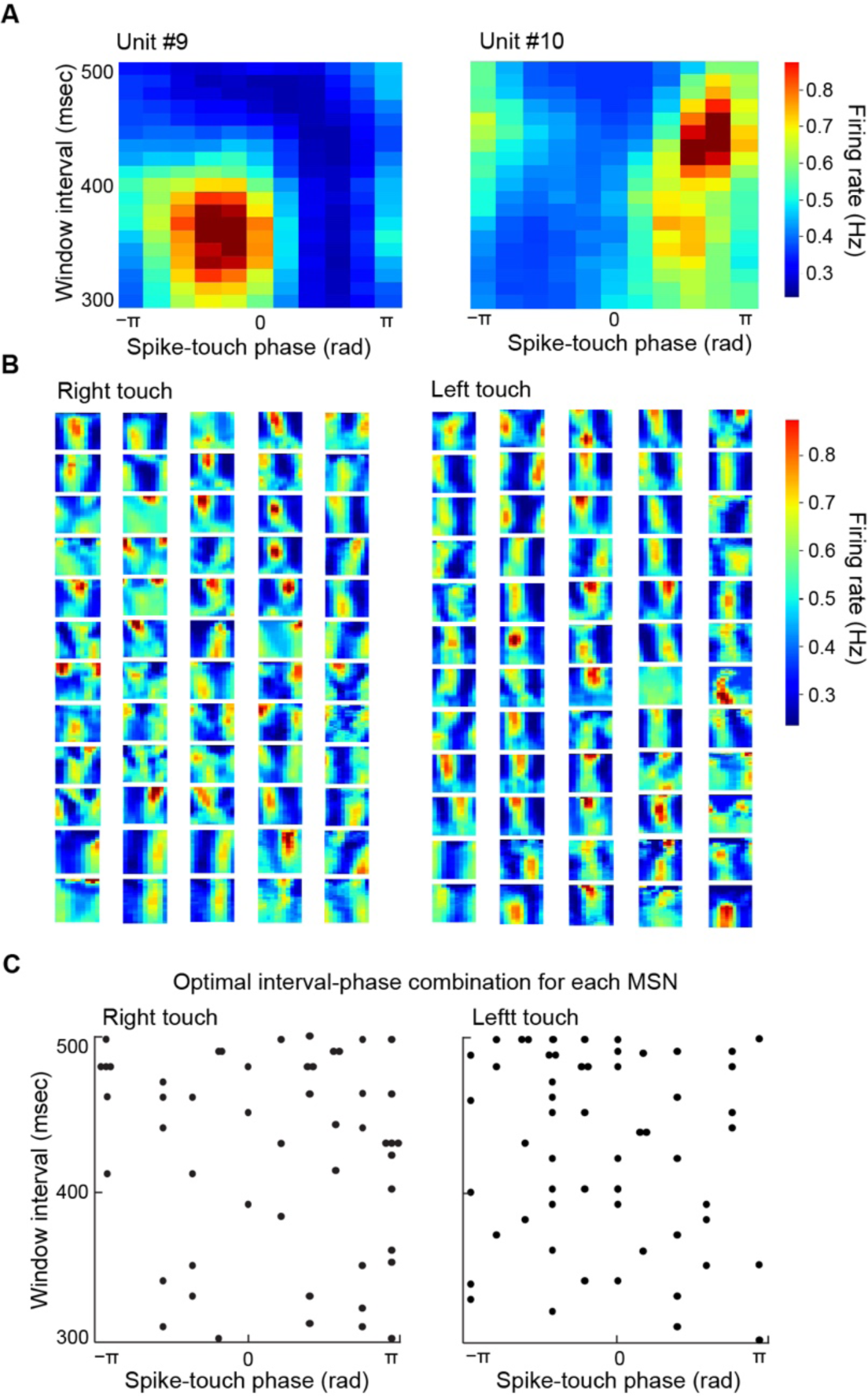
Repeat-responsive MSNs responded to optimal interval-phase combination. A. Heatmaps of interval-phase firing characteristics of two MSNs (units #9 and #10), calculated by window sets with various combinations of intervals and phases. The firing rate at the high repetition state (with 4 or more repetitive touch cycles) is shown in heatmap for each combination of interval and phase. B. Heatmaps of interval-phase firing characteristics of other repeat-responsive MSNs. See also Fig. S5 for low repetition state. C. Distribution of the optimal combination of interval and phase for each repeat-responsive MSN.

We compared the firing rates according to the number of repetitions at the optimal interval and phase for the MSNs, which was determined from the interval-phase firing characteristics maps. The firing rate of the MSNs at their optimal intervals and phases increased as the number of repetitions increased (Fig. S5B). These results clearly demonstrate that the interval responsiveness exhibited by repeat-responsive MSNs depends on multiple repetitions of a given touch interval. An interval, when repeated, is more accurately referred to as a cycle. Therefore, the interval responsiveness exhibited by the interval-responsive MSNs should correctly be considered as cycle responsiveness.

In addition, we calculated the optimal combination of interval and phase inducing a maximum firing rate for each repeat-responsive-MSN. We found that optimal interval-phase combination for each repeat-responsive MSN covered the entire range of interval and phase (Fig. 6C). This result indicated that repeat-responsive MSNs form rhythm receptive fields that represent the interval-phase combination of the forelimb movements.

### Touch-responsive MSNs also responded to the phase within the licking cycle

We recorded licking, another repetitive movement, in addition to left and right touches of the forelimbs. We have found that the relationship between left and right touches was represented as the phase by the phase-responsive MSNs. We accordingly asked whether licking and touching could both be related as phases by the MSNs, even though licking exhibits very different intervals from those of the touches.

We found MSNs that fired at a specific phase within licking cycle (Fig. 7A). Those MSNs that exhibited significant increases of firing rates at a particular phase within the licking cycle were identified as a lick-phase-responsive MSN (n = 33 units; see details in STAR Methods). We also found that the optimal firing phase within the licking cycle was different for each lick-phase-responsive MSN, covering the entire licking cycle, although approximately half of the MSNs fired around the timing of licking (0 or 2π). Further, we found cross-phase-responsive MSNs, which responded simultaneously to touch and licking phases (n = 7 units; Fig. 7C). These cross-phase-responsive MSNs fired depending on the phases of three body part cycles: left touch, right touch and licking (Fig. 7D). We identified these cross-phase-responsive MSNs as units that increased their firing rates significantly at a particular phase within the left or right touch cycle and licking cycle (see STAR Methods for details).

**Figure 7.**
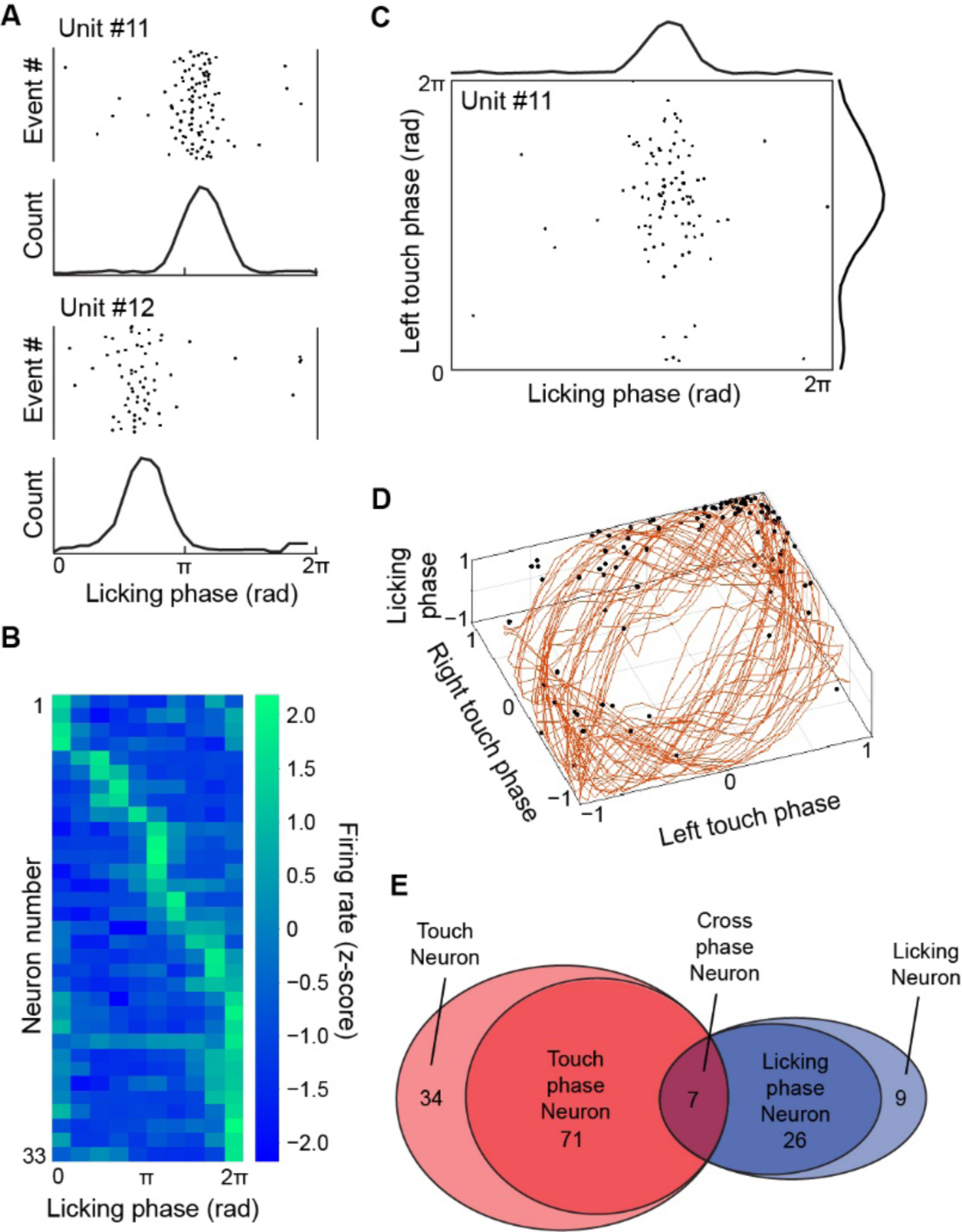
Striatal MSNs responded to combination of phases within multiple body part movements. A. Raster plots and histograms of firing events of two striatal MSNs (units #11 and #12) plotted in relation to licking cycles. B. Heatmap of average firing rates within licking cycles for all lick-responsive MSNs, sorted based on the position of their peak firing rates. C. The distribution of firing events of a single MSN (unit # 11) relative to the phase within the cycles of the left peg touch and licking. The spike histograms for each axis are shown at top and right. D. Three-dimensional plot of firing events relative to the phase within the cycle of the left touch, right touch and licking. The phases of the left touch, right touch and licking cycles are scaled to a range of −1 to 1 using a cosine function and depicted as orange lines at intervals of 10 msec, providing a visual representation of the mouse’s movement trajectory in phase space. E. Venn diagram showing the number of identified touch-responsive, touch phase-responsive, lick-responsive, lick phase-responsive and cross-phase-responsive MSNs.

In summary, we found phase-responsive MSNs for touches (n = 78 units), phase-responsive MSNs for licking (n = 33 units), and cross-phase-responsive MSNs (n = 7 units) among the touch- or lick-responsive MSNs (Fig. 7E). These findings for MSNs accord with our observation that, during running on the complex peg-pattern, motor chunks, defined as stable series of movements, are formed by appropriately coordinating the rhythm of touching and licking^8^. These phase-responsive MSNs may contribute to this coordination of multiple body parts.

### Cortical touch-responsive neurons were rarely responsive to multiple rhythm parameters

To explore whether there are neurons in the cortex exhibiting similar activity patterns as the MSNs described above, we performed recordings from cortical regions including M1, S1, and sensorimotor cortex (Fig. 8A–C). The recordings were primarily performed in the forelimb regions of the sensorimotor cortex (Fig. 8A). This specific cortical region was chosen because of its well-known extensive projections to the dorsolateral striatum^30, 31^, which included the recording sites in this study. Due to the limited number of neurons recorded from each cortical region, we analyzed all neurons collectively, pooling data from these cortical regions.

**Figure 8.**
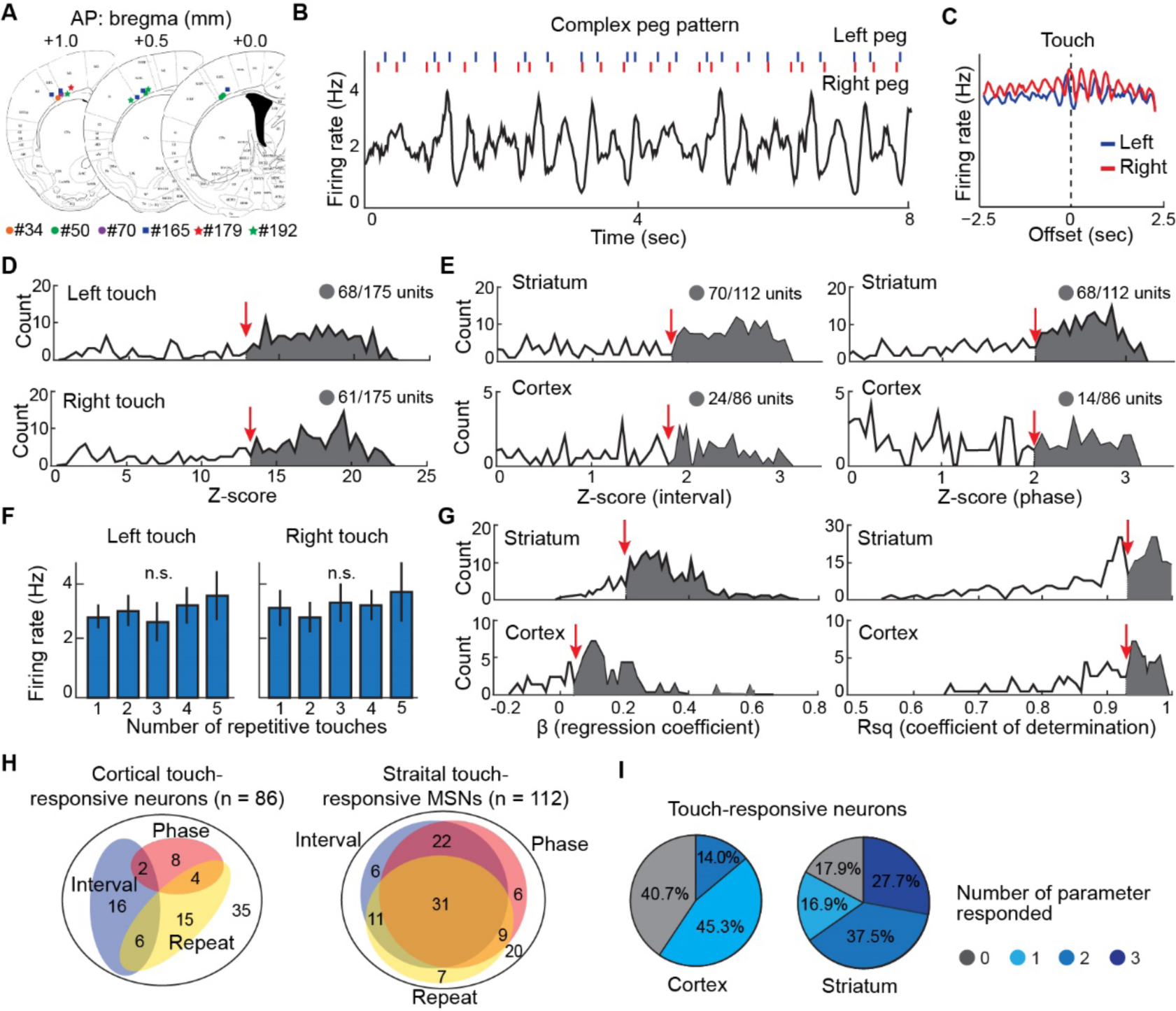
Comparison of responsiveness between cortical neurons and striatal MSNs. A. Cortical recording sites. Dots show the final locations of tetrode tips, with colors coding for individual mice. B. Firing rate of a single neuron (bottom) recorded in region of the sensorimotor cortex of a mouse running on a complex peg pattern (top), aligned to the turn marker timing. C. Cortical neural activity aligned to the time of left (blue) and right (red) forelimb peg touches. D. Distributions of peak z-scored power of the spectrogram calculated from cross-correlogram of spikes aligned to left (top) and right (bottom) peg touches. These distributions serve as the criteria for identifying touch-responsive cortical neurons. The thresholds for z-scored power are indicated by red arrows, and neurons exceeding the threshold are shown with gray shading. E. Distributions of peak z-scored power of the spectrogram calculated from cross-correlogram of spikes aligned to the turn marker, which serve as the criteria for identifying interval-responsive (left) and phase-responsive (right) neurons in the striatum (top) and the cortex (bottom). F. Analysis of repeat responsiveness in the touch-responsive cortical neurons. There was no significant increase in the firing rate of cortical neurons with respect to the number of repetitions (p > 0.05, determined by one-factor ANOVA followed by Bonferroni test). G. Distributions of the regression coefficient (β, left) and coefficient of determination (Rsq, right) as a criterion for repeat-responsive neuron. Linear regression analysis was performed on data normalized to set the mean firing rates to be 1 Hz for each neuron. H. Venn diagram illustrating the population of interval-responsive (blue), phase-responsive (red) and repeat-responsive (yellow) neurons among touch-responsive neurons in the cortex (left) and striatum (right). I. Proportion of the neurons responded to 0–4 rhythm parameters in the striatum (left) and cortex (right). Colors indicate the number of rhythm parameters.

The analyses demonstrated the presence of a distinct population of touch-responsive neurons in the neocortex (Fig. 8D). However, the number of interval- or phase-responsive neurons was relatively small (Fig. 8E, H). In addition, we could not find a consistent population-level increase in response to the number of repetitions in the cortical neurons (Fig. 8F). Repeat-responsive neurons in the cortex were defined using the same criteria as in the striatum (Fig. 8G), and interval-, phase-, and repeat-responsive neurons in the neocortex were identified (Fig. 8H). In striking contrast to the striatum, however, the sensorimotor cortex exhibited only a small number of neurons that were responsive to multiple rhythm parameters (Fig. 8I). Whereas 62.4% (n = 53/85 units) of interval- or phase-responsive MSNs in the striatum responded to both interval and phase, 5.6% (n = 2/36 units) of interval- or phase-responsive cortical neurons responded to both parameters. Furthermore, whereas 27.7% (n = 31/112 units) of touch-responsive MSNs in the striatum were responsive simultaneously to all three rhythm parameters, none (n = 0/86 units) of the touch-responsive cortical neurons displayed such responsiveness.

## Discussion

Our findings strongly argue that encoding of rhythm parameters occurs in populations of striatal MSNs. We demonstrate the presence of an extensive set of striatal neurons identified as MSNs that have selective responses to the rhythm parameters of reinforced running movements, including interval, phase and repetition of stepping and licking. To examine these features, we created a movement paradigm by use of the step-wheel that allowed experimental control of these parameters as mice ran on the wheel to receive liquid reward. We found frequent overlapping of the responses to these different parameters in many of the MSNs recorded, which likely form a tightly coupled class of MSNs. By introducing irregular patterns for the pegs that the mice had to learn to touch in order to progress at the fixed turning speed of the step-wheel, we were able to ask whether the rhythmic groupings of neural responses of the MSNs were built during stepping and licking together. We found that there was higher firing of the MSN within these chunked behavioral groupings than outside of them. We suggest that the formation of these condensed periods of firing and associated stepping and licking behaviors represented the process of chunking, a well-known property of movement organization^32–34^ proposed as one function of neurons in the sensorimotor striatum^35^.

We first identified MSNs that responded to forelimb touches on the pegs (135/349 units). It was among these that we identified MSNs that were activated depending on the intervals, phases, and the numbers of repetitions of the left and right forelimb touches. Many of these MSNs possessed firing rates that were sensitive to optimal intervals of forelimb touching. The peak interval responses of these MSNs covered almost the entire range of the intervals that occurred in the interval-changing peg-pattern. Many of the touch-responsive MSNs also possessed optimal left-right differences in forelimb position. The model analysis of predicting neural activity from touch timing suggested that this optimal left-right difference might be produced by the superimposition of relative, rather than absolute, timing within a single cycle of each left and right limb movement. The relative timing in a cycle can be expressed as the phase. The results thus indicate that touch-responsive MSNs used phase to represent the coordination between forelimb movements. We also found striatal MSNs that responded to both touch and lick phases, suggesting that these MSNs had encoded the phase between movements of different cycles. These findings indicate that the striatal MSNs could connect multiple body movements of different frequencies by multiplexed encoding the phase information between each body part movement.

In addition to interval and phase responsiveness, the firing rate of touch-responsive MSNs increased as the numbers of touches were cyclically repeated. This was a crucial finding. When a particular interval is repeated, a rhythm arises, and the interval of the movement can be taken as a cycle. Thus, for many of the touch-responsive MSNs that changed their firing rates in response to intervals and the number of repetitions, their interval responsiveness can be regarded as cycle responsiveness. Given that the phase represents a relative position within a cycle, we suggest that these MSNs could represent by their ensemble firing a combination of continuous movements by the phase during which the movements are highly repetitive, which means rhythmic encoding. We refer to these touch-responsive MSNs collectively as the rhythm-responsive MSNs. Considering the distinctive responsiveness of individual rhythm-responsive MSNs and their collective covering of the entire representation within the interval-phase dimension, it becomes plausible to conceptualize that these MSNs have *rhythm receptive field*s.

Combinatorial use of striatal rhythm-responsive neurons forming the rhythm receptive fields may allow whole-body movements to be represented as rhythmic combinations, perhaps by selecting an appropriate set of downstream central pattern generators. Increasing evidence suggests that small groups of brainstem neuron encode highly discrete motives of movements (for example, moving forward but not other movements)^36, 37^. It has already been found that the pars reticulata of the substantia nigra, a key basal ganglia output nucleus, provides input to some of these^38^. The combinatorial encoding that we have found in the MSN rhythm receptive fields may provide a means for integrating distinct motives into purposeful continuous movements. It is known that the striatum, in its role of controlling movement, properly regulates downstream direct-indirect pathways^39–41^. We did not identify the MSNs recorded as direct or indirect spiny projection neurons, and we did not observe obvious groupings within the MSN ensemble that responsive to rhythm. It is thus a challenge to understand how these neurons would drive the striatal output regions of the pallidum and substantia nigra through its well-known D1 and D2 receptor-mediated direct and indirect pathways^39–41^. These could have opposing or cooperative actions, depending on the requirements of the task types, promoting or regulating movement sequences. Further work could address their specific impact on these circuit-based movement controls with combinations of cell-specific tagging and recording.

One of the main excitatory inputs to the striatum arrives from the cerebral cortex. The dorsolateral striatum from which neurons were recorded in this study receives inputs from sensory and motor areas of the cerebral cortex^30, 31^. We found that a majority of striatal touch-responsive MSNs were responsive to more than two rhythm parameters concurrently, whereas cortical touch-sensitive neurons were few in percentage and only responsive to one or two. These findings indicate that the striatum may have a key function in integrating cortical inputs that separately encode single or rarely two rhythm parameters. Moreover, the presence of cross-phase MSNs in the striatum suggests that the striatum also integrates rhythm information of multiple body parts. Thus, our findings suggest that the striatum encodes and integrates the rhythm information derived from diverse inputs originating from different sensorimotor areas of the cortex, and consequently subserves movement coordination of multiple body parts.

The striatum also receives the dopamine-containing input from the substantia nigra pars compacta, which is strongly impacted in PD. Disabling movement problems occur, including bradykinesia, disabled gait and akinesia^42, 43^. Notably, loss of motor rhythm has been reported in patients with PD^25, 26^. In addition, in patients with PD, rhythmic cues through visual or other modality improve motor function such as walking^44^, and are used as an effective rehabilitation method. Collectively, these studies suggest that PD patients may have impaired ability to establish rhythmic coordination of movement, which should be constructed in the brain during exercise or motor planning. Given that the striatum is the primary projection site of dopamine, it is plausible that this rhythmic coordination ability may reside in the striatum, which aligns with the results of this study.

We have shown in a behavioral study employing the step-wheel that the dorsal striatum plays an important role in walking and running in the wheel, and that inhibition of dopamine receptors in the dorsal striatum significantly reduces their speed of running^11^. These results, together with the findings reported here, have direct relevance to the gait abnormalities and impaired coordination observed in PD patients during walking. It is possible that some of these difficulties stem from the malfunction of rhythm receptive fields provided by the striatal rhythm-responsive neurons in processing rhythmic information. The weak synchronization between limb movements and other accompanying movements in the gait of PD patients accords with this hypothesis. Much research also suggests that the striatum is involved in the control of movement speed and vigor^15–18^. Considering that lack of movement coordination can lead to decreased movement speed and vigor, our findings support this idea. PD patients further have deficits in motor initiation and termination. In addition, as several studies have demonstrated that neurons in the striatum are required for the initiation and termination of sequential actions^19, 45–47^, it is reasonable to think that the activity of the MSNs responsive to ‘approach’ and ‘leave’ found here could contribute to these striking symptoms of PD.

It has been noted that computing coordination among multiple body parts separately would demand unwieldy levels of computation and make the computation of execution inefficient because of the infinite number of possible combinations of movements^1^. But if body parts are interrelated in movement sequences, the computational load can be significantly reduced. For continuous repetitive movements, the use of rhythm parameters for interrelationships between body parts seems to be an efficient method. We found that MSNs in the sensorimotor sector of the striatum responded selectively to various cycle lengths and repetitions and that forelimb relationships were represented as phases. These findings demonstrate that rhythm-based representation in the striatal MSNs by rhythm receptive fields could be used for continuous movement execution and even for combinations of movements of different cycle lengths. This suggests the potential integration of multiple body parts into fewer rhythms, resulting in dramatic savings in computational load. This saving is probably feasible due to the fact that many pairs of movements, such as breathing and walking, are integer multiples of frequency^48–50^.

We found many MSNs that exhibit increased firing rates in response to specific combinations of movement intervals and phases, marking a novel dimension of neural specialization. In contrast to classical visual receptive fields in the visual cortex^51^, or place fields of the hippocampus, which process spatial and mnemonic features^52, 53^, these MSNs respond to temporal patterns of movement, aligning with the striatum’s role in motor control and coordination. The brain, as here represented by the striatum, may use these rhythm parameters to improve efficiency by representing complex sequential movements as a combination of rhythms. Spinal and brainstem mechanisms are themselves attuned to coordinating encoding features as well^54, 55^, so that one can perhaps best view the coordination of rhythmic features found here for the striatum as supraordinate frameworks guided by corticothalamic inputs.

## Acknowledgements

We thank Dr. Daniel J. Gibson for his critical reading of the manuscript and excellent suggestions for the analysis. This research was supported by the JST SPRING (JPMJSP2138), the Nakatomi Foundation, the National Institute of Mental Health (R01 MH060379), and the KAKENHI (18H04945 and 22K18662).

## Author contributions

Conceptualization, Y.K., A.M.G., and T.K.; Methodology, T.N., H.D., Y.K., A.M.G., and T.K.; Software, K.H., T.N., T.T., and T.K.; Formal Analysis, K.H., T.N., T.T., and T.K.; Experimental work, K.H., T.N., T.T., H.D., Y.K., and T.K.; Writing – Original Draft, K.H.; Writing –Review, further analyses, and concept formation, K.H, A.M.G, and T.K.; Visualization, K.H., T.N., T.T., Y.K., A.M.G., and T.K.; Supervision, T.Y., A.M.G., and T.K.; Funding Acquisition, K.H., T.Y., A.M.G., and T.K.

## STAR Methods

### RESOURCE AVAILABILITY

#### Lead Contact

Further information and requests for resources and reagents should be directed to and will be fulfilled by the Lead Contact, Takashi Kitsukawa.

#### Data and code availability

- All data reported in this paper will be shared by the Lead Contact upon request.
- All original codes used in this study have been deposited at Github and are publicly available as of the date of publication.

### METHOD DETAILS

#### Animal

ICR mice (10–20 weeks old, n = 15) were used in the experiment. Mice were kept in home cages with free access to water and food, and were water-restricted from one day before starting the training session on the step-wheel task until the end of the experiment. Water was given only while performing the step-wheel task during the water restriction period, but they were given free access to water for an entire day every 1–2 weeks. Their health condition, including weight, body temperature and fur condition, was closely monitored during water restriction. Water was given ad lib if their weight fell to less than 80% of its initial level or if any abnormal health condition was observed. All procedures for the following experiments were approved by Osaka University animal experiments committee and performed according to the guidelines for animal experiments of the Osaka University.

#### Step-wheel

The step-wheel consists of two overlapping motor-driven rotating disks (Fig. 1A, Kitsukawa et al.^9^). The disks have a diameter of 32 cm, and a circumference of 100 cm. Pegs (25 mm) are attached to the left and right disks to serve as footholds for the mice. The experimenter can change their placement, thus creating a series of peg-patterns^7, 9^.

Four types of peg-patterns were used in our study: regular, complex, interval-changing and phase-changing peg-patterns (Fig. 1B). In the regular peg-pattern, the left and right pegs were alternately and equally spaced, whereas in the complex peg-pattern, the pegs were placed in an irregular arrangement with no specific rules. In the interval-changing and phase-changing peg-patterns, peg spacing and left-right relative position were changed gradually within the pattern, respectively. Although the left and right pegs were arranged irregularly in the complex peg-pattern, the total number of pegs arranged in a circle was the same as in the regular peg-pattern. Thus, the average peg-to-peg spacing was the same between the regular and complex peg-patterns. Because one trial was defined as one peg-pattern that spanned over a one-half round of the wheel, one round of the wheel consisted of two trials. There was a water spout between the left and right pegs, from which the mouse could receive water as a reward. Because the wheel was rotating, the mouse was required to run on the pegs to obtain water.

In addition, voltage sensors were attached to each peg and drinking spout to detect the timing of limb touches to the peg and licking of the spout^9^. Infrared sensors were also placed to detect when the mouse was in the vicinity of the drinking spout. In this study, we analyzed the neural activity only when the mice blocked this infrared light.

#### Pre-training

On the pre-training session day 1, water was supplied to the mice inside the step-wheel for 10 min without rotating the wheel to allow the mice to learn the location of the water spout and adapt to the environment. On the pre-training session day 2, the mice initially ran at a speed of 20 sec per trial (25 mm/sec) on the regular peg-pattern. The rotational speed gradually increased over the next 5 days to 1 week, and the peg-pattern was changed to the complex peg-pattern. Pre-training was completed when the mice could run at a speed of 4 sec per trial (125 mm/sec) on the complex peg-pattern. Mice were then operated for the implantation of head stage (see below). Water restriction resumed after a one-week post-operative recovery period, and pre-training was restarted. Recovery sessions after the surgery were completed in approximately one week.

#### Head stage

A head stage with eight built-in tetrodes was prepared to record neural activity from the dorsolateral striatum of running mice. The tetrodes were made by bundling four 10-µm diameter nichrome wires^56, 57^. The head stage used in this study was manufactured by Neuralynx. DiI was applied to the tip of the tetrode to locate the recording site in the brain.

#### Surgery

After the pre-training sessions, a surgical procedure was performed to fit the head stage. Mice were anesthetized by injecting mixture of three anesthetics (0.3 ml of Domitor (medetomidine), 0.8 ml of Dormicum (midazolam), and 1 ml of Betorfal (butorphanol) in 19.75 ml of saline) at the dose of 10 µl/g of mouse body weight. Ten min after the anesthesia was administered, we confirmed that the anesthesia had fully taken effect and made incision to expose the skull. The height of the bregma and lambda was adjusted horizontally. A hole was drilled in the skull (AP: 0.5–1.0 mm, ML: 2.5–3.0 mm), and the dura mater was removed. The head stage was fixed to the brain surface with dental cement. After one week of recovery period after surgery, the water supply was again restricted, and training was resumed. During the post-operative recovery period and training, the tetrodes were gradually advanced by rotating a screw attached to the head drive (approximately 250 µm per rotation) and lowered approximately 1–1.2 mm from the brain surface. At that time, we confirmed that the tip of the tetrode reached the dorsolateral striatum by observing neural activity. Specifically, we determined that the tetrode reached the dorsal striatum when it passed the white matter layer, where typical cortical neural activity was lost, and very little activity was observed, and then approached a region that showed phasic burst firing while running, although spontaneous firing rate was low^58^.

#### Recordings

A 32-channel preamplifier (Neuralynx, MT) was attached to a circuit board mounted on the head stage during striatal neural activity recording from mice. The obtained data were first amplified by preamplifiers attached to the head stage, sent to an 8-channel amplifier (gain: 10,000, filter: 0.6–6 kHz) and stored by the Cheetah Data Acquisition System (Neuralynx). Spike sorting was performed manually using SpikeSort3D (Neuralynx). Cluster separation was performed mainly by analyzing peak values of spike amplitudes, but sometimes energy and valley of spike amplitudes were used as references to improve accuracy. Clusters with spikes with inter-spike intervals less than 1 msec were excluded from the analysis. Spike data were analyzed with custom software created using MATLAB (MathWorks, MA).

#### Behavioral data

Behavioral data of running mice were recorded using voltage sensors and infrared sensors attached to the step-wheel. The timing of when the mice touched the pegs and put their mouths to the water spout (licking) was detected by recording the electrical potentials applied to the pegs and water spout using LabVIEW (National Instruments, TX), which were connected to a 20-kHz oscillator through separate resistors to monitor contacts with the grounded animal. The timing of mice approaching and leaving the water spout was detected using an infrared sensor placed above the water spout. Data obtained from the experiments were analyzed using custom software created in MATLAB and Python.

#### Recording session

One week after surgery, recovery training resumed with the head stage connected to the recording cable. The training was continued until the mice were able to run one session at a speed of 4 sec per trial. The following recording experiments were then performed. On the day of recording session, 13 mice ran on three types (interval-changing, phase-changing and complex) of peg-pattern. However, the remaining 2 mice only ran on the interval-changing and phase-changing peg-patterns. Mice were allowed to run each peg-pattern for 30–60 trials at a constant speed (3.5– 4.5 sec per trial). After recording the neural activity on each peg-pattern, we advanced the tetrode slightly to search for other neurons, and the recording session was repeated on the following day, until the tetrode position reached approximately 1.8 mm below the brain surface.

#### Recording site location

The electric current was applied to mark the final tetrode tip locations after the completion of data collection, and 24 hours later, the brain was perfusion-fixed with 4% paraformaldehyde under anesthesia with an excess volume of the anesthetic solution described above, and the brain was removed. The removed brains were trimmed and post-fixed in the same fixative for 24 hours and then replaced with 30% sucrose. Then, 30-µm sections were prepared using a freezing and sliding microtome and placed on glass slides. After sections were dried, Nissl staining was performed, and the location of tetrodes was confirmed under a microscope.

#### Classification of striatal neuron types

Units identified by spike sorting using SpikeSort3D (Neuralynx) were classified into medium spiny projection neurons (MSNs), fast-spiking interneurons (FSIs) and other neurons. To distinguish between different types of neurons, peak-to-trough and half-peak widths were calculated from the spike waveforms, and clustering was performed based on these waveform features, following the methods described in previous studies^27–29^. Neurons with a peak-to-trough width greater than 0.31 and a half-peak width greater than 0.16 were determined as MSNs, and those with a peak-to-trough width less than 0.31 and a half-peak width less than 0.16 were determined as FSIs. Units with a low firing rate (< 0.5 spikes/sec) and those that had short (<1 msec) inter-spike intervals in more than 0.5% of all inter-spike intervals were excluded from the analysis. Given that the step-wheel equipped voltage sensors attached to all the pegs and spouts, recorded data contained relatively large amount of noise throughout the session. Therefore, we set a strict threshold to better ensure that we could correctly identify MSNs, which resulted in a relatively lower rate of MSN identification.

#### Relationship between movement and neural activities

The neural activities of all mice (n = 15) were included in the analysis. Cross-correlograms were generated to examine the temporal relationship between the neural activity and the timing of peg touch, licking, and approach to/leave from the water spout. For the cross-correlogram related to peg touch, we calculated the firing rate by aligning the neural activity to the timing of peg touch (bin: 10 msec, moving average: 100 msec, range: ±2.5 sec around the event). In the cross-correlogram for licking, neural activity was aligned by the timing of licking (bin: 4 msec, moving average: 20 msec, range: ±1 sec around the event). For the cross-correlograms related to approach to/leave from the water spout, neural activity was aligned by the timing when the mouse blocked/released the infrared sensor (bin: 20 msec, moving average: 200 msec, range: ±5 sec around the event).

#### Classification of touch- and lick-responsive neurons

We applied criteria to classify neurons that responded to peg touch or licking based on cross-correlograms. Neurons with an average firing rate of less than 0.5 Hz during the one-day running session were excluded from the analysis.

To identify striatal MSNs that responded to peg touch, we first performed a DFT on the cross-correlograms of the spikes aligned by left and right peg touches and obtained a spectrogram. The range of offsets computed in the cross-correlograms were set to 5 sec (−2.5 to 2.5 sec). The spectrogram was then converted to z-scores, and z-scored power was calculated. In the step-wheel running, the average interval between two consecutive peg touches was 370 msec (2.7 Hz) despite the diverse left-right limb positioning. Then, we considered a z-scored power of more than 17.5 for left touch and more than 17 for right touch in the 2–3.5 Hz frequency band as indication of responsiveness to left and right peg touch, respectively (Fig. S1A, see details below). Neurons that were responsive to either left or right peg touch were then defined as touch-responsive neurons (Fig. 2D).

Similarly, for neurons that responded to licking, we performed a DFT on the cross-correlogram of spikes aligned by licking. The range of offsets computed in the cross-correlograms were set to 2 sec (−1 to 1 sec). Neurons with a z-scored power of more than 11.5 in the 7–11 Hz frequency band in the spectrogram were considered responsive to licking (Fig. 2E and Fig. S1B, see details below), because the average interval between licking was 110 msec (9.1 Hz).

For the cortical neurons, touch-responsive neurons were determined using the same method as in the striatal MSNs. A threshold was set based on the distribution of z-scored power in the spectrogram (Fig. 8D). Specifically, z-scored power of more than 12.5 for left touch and more than 13 for right touch was used as criterion for responsiveness to left and right peg touch, respectively. Z-scored power was calculated from the Fourier transformed spectrogram in the frequency band of 0–10 Hz for peg touch and in the frequency band of 0–25 Hz for licking. Each neuron had neural data from more than two sessions with non-regular peg-patterns, and the response property of each neuron was determined based on the neural data with the highest z-scored power for the neuron. Then, the distribution of z-scored power for all individual recorded neurons was plotted (e.g., Fig. S1A). To determine the threshold of z-scored power for detecting touch- and lick-responsive neurons, we manually set the threshold at the point of upward deflection in the distribution curve, which effectively separated the group of neurons with high z-scored power.

#### Classification of approach- and leave-responsive neurons

To identify neurons that responded to the approach to/leave from the water spout, we used data obtained for all peg-patterns and compared the firing rate during ±1-sec period around the timing of infrared blockade or release, respectively, We performed two-tailed t-test on spike data between those two periods, then the neurons with significantly higher (p < 0.05) firing rates before the blockade were determined as approach-responsive neurons, and those with higher firing rate after release were determined as leave-responsive neurons.

#### Interval-changing and phase-changing peg-patterns

The interval-changing peg-pattern refers to a peg-pattern where the spatial interval between the pegs gradually widens and then returns to a narrower interval within a single trial. In this pattern, the positional relationship between the left and right pegs remains fixed (face-to-face) to minimize the influence of positional differences between the left and right limbs. The interval is approximately doubled during the first half of the trial and then returns to the initial interval in the second half.

The phase-changing peg-pattern, on the other hand, maintains a constant and unchanging peg interval on each side, but the interval differs between the left and right sides. This pattern includes a gradual change in the relative positions of the left and right pegs. Initially, the left and right peg positions start in a face-to-face state. Then, in the first half of the trial, the left and right peg positions become alternate, and in the second half, they return to the initial state.

#### Classification of interval- and phase-responsive neurons

Criteria were established to classify neurons that respond to interval and left-right differences. Considering that one rotation of the step-wheel consists of two trials, the peak firing rate of MSNs responsive to interval or left-right differences occurred twice within one wheel rotation.

To identify striatal MSNs that responded to interval and phase, we first performed a DFT on the cross-correlograms of the spikes aligned by the timing of trial start and obtained a spectrogram. The range of offsets computed in the cross-correlograms were set to 8 sec (0 to 8 sec). For the identification of interval-responsive MSNs in the striatum, touch-responsive MSNs with z-scored power of 1.8 or higher at 0.25 Hz frequency in the spectrogram during running on the interval-changing peg-pattern were identified as interval-responsive MSNs (Fig. 8E). Similarly, for phase-responsive MSNs, touch-responsive MSNs with z-scored power of 2.0 or higher at 0.25 Hz frequency during running on the phase-changing peg-pattern were identified as phase-responsive MSNs (Fig. 8E).

In the cortex, the same distribution-based approach was used to determine the threshold for identifying interval- and phase-responsive neurons. As there were populations with high z-scored power for each interval and phase in striatal touch-responsive MSNs, a manual threshold was set to effectively discriminate between these populations. By contrast, due to the absence of clear populations with high z-scored power for both interval and phase among cortical touch-responsive neurons, the threshold determined for MSNs was applied to cortical data to define interval- and phase-responsive neurons in the cortex.

To determine the location of the maximum envelope in each neuron, the low-frequency information was extracted by applying the low-pass filter to the firing rates for one round of the step-wheel. Then, histograms were z-scored for each round. This process was applied to the firing rates of interval- or phase-responsive MSNs to generate heatmaps shown in Fig. 3E and 3F. The MSNs shown in the heatmaps were sorted and rearranged from top to bottom based on the peak position of the firing rate within a trial.

Phases were fixed to zero (face-to face peg arrangement) in the interval-changing peg-pattern, which means that we evaluated interval-responsiveness only for neurons that responded even slightly to the face-to-face pattern. Similarly, phase-responsive neurons that we identified responded to the intervals used in the phase-changing peg-pattern. However, it is not realistic to use a large number of peg-pattern variations covering entire interval and phase combinations. Thus, in this experiment, we used these two patterns to identify the interval- and phase-responsive neurons.

#### Decoding analysis of neural activity in phase-responsive neurons

To test the hypotheses 1 to 6 models 1 to 6 were constructed to predict the firing rate using the peg touch data collected during running on the complex peg-pattern. The model aimed to capture the relationship between the peg touch pattern and the firing rate of neurons. Subsequently, the firing rate was predicted from the peg touch data collected during running on the phase-changing peg-pattern. The predicted firing rate was then compared with the actual firing rate of the neurons recorded during the phase-changing peg-pattern (Fig. S4). Specifically, the following calculations were performed for each model.

##### Model 1

The firing rate of neurons within one cycle was calculated based on the phase in the cycle (relative time, bin = 0.05 cycle). Based on these data, models were created separately for each left and right limb to predict the firing rate with “relative time” within one cycle as input to predict the firing rate of neurons.

##### Model 2

The firing rate of neurons within one cycle was calculated based on the time interval between neural activity and subsequent touch (absolute fire-to-touch time, bin = 20 msec). Separate models were created for each left and right limb, using the “absolute fire-to-touch time” within one cycle as input to predict the firing rate of neurons.

##### Model 3

The firing rate of neurons within one cycle was calculated based on the time interval between neural activity and preceding touch (absolute touch-to-fire time, bin = 20 msec). Separate models were created for each left and right limb, using the “absolute touch-to-fire time” within one cycle as input to predict the firing rate of neurons.

For models 1 to 3, when predicting the firing rate in the phase-changing peg-pattern, the actual peg touch data were used as an input to predict the firing rate using the left and right limb models separately. The firing rate of neurons in one session was then predicted by calculating the weighted sum of the left and right model outputs, taking into account of the neural responsiveness to the left and right touches. The predicted firing rate was averaged within a wheel rotation (2 trials), smoothed with a 50-msec window, and finally compared with the actual firing rate during running on the phase-changing peg-pattern.

##### Model 4

The firing rate of neurons inside a left touch cycle was calculated according to the time interval between the right touch and the subsequent left touch (absolute right-to-left touch time, bin = 20 msec). Based on those data, a model for predicting the firing rate was created with “absolute right-to-left touch time” as the input.

##### Model 5

The firing rate of neurons inside a left touch cycle was calculated according to the time interval between the right touch and the preceding left touch (absolute left-to-right touch time, bin = 20 msec). Based on those data, a model for predicting the firing rate was created with “absolute left-to-right touch time” as the input.

##### Model 6

The firing rate of neurons inside a left touch cycle was calculated according to the right touch phase within the left touch cycle (relative time, bin = 0.05 cycle). Based on those data, a model for predicting the firing rate was created with “relative time of right touch within the left touch” as the input.

For each of these models (4, 5, and 6), the predicted firing rate was averaged in a wheel rotation (2 trials), smoothed with a 50-msec window, and finally compared with the actual firing rate during running on the phase-changing peg-pattern.

#### Evaluation indices for the model

To evaluate the prediction accuracy of the firing rate for phase-responsive neurons during running on the phase-changing peg-pattern, two evaluation indices, peak error and JSD, were used.

Peak error measures the x-axis distance (time direction) between the maximum local value of the predicted and actual firing rates around each touch. The peak error was calculated by comparing the closest peaks and by averaging the calculated distances. JSD was used as another evaluation index. It assesses the similarity between the predicted and actual distributions of the firing rate within one rotation of the wheel. The local maximum values of the firing rate were extracted, and the envelopes connecting those local maxima were obtained. The firing rate was then converted into a probability distribution by normalizing the area under the envelope to be 1. Finally, JSD was calculated between the predicted and actual distributions to obtain the error index.

The prediction accuracy was evaluated using these two indices for phase-responsive MSNs (n = 68 units) recorded from 13 mice that ran the complex peg-pattern and phase-changing peg-pattern. To evaluate the differences in prediction accuracy among the hypotheses, one-factor ANOVA followed by Bonferroni test was performed.

#### Phase responsiveness within one cycle

To generate a histogram of neural activities for striatal MSNs within one cycle, the spikes were counted throughout the session. The duration of one cycle was divided into 20 bins. For each neuron, the number of spikes corresponding to the phase within the cycle was calculated separately for each left and right touch cycle.

#### Sorted heatmap of the phase responsiveness within one cycle

The number of spikes within one cycle was calculated and subsequently z-scored within each session. Furthermore, the location within the cycle where the spikes exhibited the highest occurrence was determined for each neuron. The histograms of spikes for all neurons were then sorted based on the peak location of spikes within the left and right cycles, respectively. These sorted histograms were represented as a heatmap, allowing for the examination of neural activity patterns across different phases of the cycle (see Fig. 4C).

#### Window analysis

Window analysis was performed on touch-responsive MSNs recorded from mice running on the phase-changing, interval-changing and complex peg-patterns (Table 2). Six windows, consisting of five touch windows for detecting peg touches and one spike window for detecting spikes, were used in this analysis. The touch windows were evenly distributed along the time axis, whereas the spike window was positioned either before or after the third (center) touch window. By sliding this window set along the time axis of the data, touch and firing events falling within the windows were counted at each time step.

The placement of touch windows at regular intervals facilitated the quantification of repetitive touch cycles (0 to 5). On the other hand, the positioning of the spike window within the touch windows allowed for the calculation of firing rates at specific phases within the cycle. Through the window analysis, it is possible to examine the firing rates of neurons in relation to the periodicity of movement rhythm.

A total of 21 types of touch windows were prepared, spaced at 300–500 msec intervals with a step size of 10 msec. To determine the positional relationship of the spike window within the touch window, the center of the touch window was assigned as 0π, the preceding touch window (one cycle before) was defined as −2π, and the subsequent touch window (one cycle after) was defined as 2π. Likewise, 11 types of spike windows were prepared, ranging from -π to π with a step size of 0.2π.

The combination of 21 touch window intervals and 11 spike window positions resulted in 231 window patterns. The subsequent analysis was performed using these patterns. The width of the touch window was set to half the interval (i.e., 150 msec if the interval was 300 msec), and the width of the spike window was set to 1/10 of the touch window interval.

Applying the window set with a sliding interval of 5 msec to the data enabled the counting of touch and firing events that fell within the windows. By dividing the firing event counts with the width of spike window, firing event counts were adjusted to firing rates. This allowed for the calculation of the number of touches within the five touch windows (indicating the repetitive touch cycle, ranging from 0–5) and the firing rate of neurons at specific phases at each time point.

#### Calculation of firing rate with repetition

During the analysis of neuronal response to repetition, a window set was created with the optimal period and phase characteristics of each neuron. The number of repetitions and firing rates were calculated by sliding the window. To determine the optimal interval for a neuron to fire, data from running sessions on the interval-changing peg-pattern were used. A bin size identical to the touch window (300–500 msec; bin = 10 msec) was used, and the firing rates dependent on touch intervals were calculated separately for left and right touches. The interval with the highest firing rate was identified as the optimal interval for each neuron.

Similarly, to determine the optimal phase, the one-cycle firing rate data from sessions with the phase-changing peg-pattern, as presented in Fig. 4C, were used. The phase within the cycle exhibiting the highest firing rate was determined as the optimal phase for each neuron. A window set, determined by the optimal interval and phase through the aforementioned method, was created.

The repetitive touch cycles and corresponding firing rates were calculated using the window analysis applied to the data obtained from running on the complex peg-pattern. Subsequently, the firing rates were averaged across the entire session according to the number of the repetitive touch cycles. To evaluate the effect of the difference in the number of repetitive touch cycles on the firing rate, a one-factor ANOVA followed by multiple comparison test of Bonferroni was performed.

#### Determination of repeat-responsive neuron

Repeat-responsive neurons were identified based on their increased firing rate in relation to the number of repetitive touch cycles. To determine the repeat responsiveness of neurons, the firing rate was calculated for each left and right limb, considering the number of touch repetitions (ranging from 1 to 5), and a window analysis was performed with the optimal interval and phase for each neuron. A linear regression analysis was then conducted to quantify the relationship between the number of repetitive cycles and the firing rate. Prior to the regression analysis, the firing rate was normalized with the average firing rate set to 1 Hz. The regression analysis was performed using the data running on the complex peg-pattern. In cases where neurons had multiple running sessions on the complex peg-pattern, the session with the most significant repeat responsiveness was selected for analysis.

To identify repeat-responsive neurons, thresholds were set at the point of deflection in the distribution curve of regression coefficients (β) for each neuron, enabling discrimination of neurons with relatively higher β values. Subsequently, the threshold for the coefficient of determination (Rsq) was set at the point of deflection in the Rsq distribution curve, allowing separation of the neuron group with high Rsq values.

For striatal MSNs, a neuron was considered repeat-responsive if it exhibited a significantly positive slope (regression coefficient β) of 0.2 or higher when analyzing the relationship between the number of repetition cycles and firing rate, along with a coefficient of determination (Rsq) of 0.93 or higher. Finally, MSNs that responded to repetition in either the left or right cycle were identified as repeat-responsive MSNs. In the case of cortical neurons, the threshold for β was set to 0.05, and the threshold for Rsq was set to 0.93, based on the distribution of these parameters.

#### Motor chunk detection

To identify chunk regions, we applied a method based on the variance of the inter-touch interval, which was previously employed in our study^8^. This approach for motor chunk detection focuses on the presence of stable and non-stable running regions observed throughout the experimental session.

Initially, for each session, the variance of the inter-touch interval was calculated for each peg. Subsequently, regions exhibiting a low variance in the inter-touch interval for four or more consecutive steps were identified as chunk regions. The firing rates of repeat-responsive MSNs within and outside these chunk regions were then calculated, followed by a two-tailed t-test to evaluate any significant differences.

#### Heatmap of interval-phase firing characteristics

The heatmap of interval-phase firing characteristics was generated through window analysis using multiple window sets. Each window set individually calculated the number of repetitive cycles and firing rate at each time. Then, the data were classified into two states: low repetition state and high repetition state. A high repetition state was defined when the number of repetitive touch cycle was 4 or more, and a low repetition state was defined when the number of repetitive touch cycle was 3 or less. For each window set (specific set of interval and phase), the average firing rates during the high and low repetition states were calculated. This calculation was performed for all interval and phase window sets, resulting in a total of 21×11 patterns, which were then presented as a heatmap (Figs. 6 and S5A).

#### Maximum firing rate for each number of repetitive touch cycle

For each neuron, a maximum firing rate was defined to measure the extent of firing rate modulation based on the number of repetitive touch cycles. Initially, the firing rate was calculated for each number of repetitions using all 21×11 window sets. The window set that yielded the highest firing rate for each number of repetitions was then selected to determine the maximum firing rate associated with that specific number of repetitions. To evaluate the difference in the maximum firing rates across various numbers of repetitions, a one-factor ANOVA followed by multiple comparisons of the Bonferroni test was performed.

#### Cross-phase analysis

Neural activity of MSNs recorded from all mice (n = 15) were used for performing cross-phase analysis between licking and left-right peg touch. A single cycle of licking was considered as phase range of 0–2π. Subsequently, all firing events occurring throughout the session were represented in a raster plot, displaying the neural activity for each individual neuron. The firing event histograms were presented in the figures below the raster plot by quantifying the firing events associated with specific phases within a single cycle of licking. The firing events were counted by dividing the licking cycle into 10 bins (bin = 0.2π). The number of firing events within each licking cycle was z-scored for individual neurons, and subsequently arranged based on the peak position of the firing event counts. These arranged firing rates were then presented as a heatmap.

For two-dimensional scatter plot, the phases within the left touch cycle and licking cycle were calculated for each firing event. Both the left touch and licking cycles were divided into 20 bins, and the firing events were counted within each bin. To ensure smoothness, the firing event counts within each bin were smoothed by averaging over ±2 bins, then they were presented alongside the scatter plot.

To mitigate the influence of the number of licks, the firing event counts were normalized by dividing the number of firing events by the number of licks in each touch phase. This normalization method effectively removed the effects of the licking count on touch phase responsiveness in each neuron. Subsequently, neurons displaying a firing event count at least twice as large as the average firing event count within a specific touch phase were identified as touch-phase-responsive neurons.

Regarding the responsiveness to lick-phases, the licking cycle was divided into 10 bins, and the firing events were counted within each bin. Neurons exhibiting a firing event count at least twice as large as the average count within a specific lick phase were identified as lick-phase-responsive neurons. Neurons exhibit responsiveness to both touch-phases and lick-phases were identified as cross-phase-responsive neurons.

To facilitate visualization, the phases of firing events within one cycle of left touch, right touch and licking were scaled to a range of −1 to 1 using a cosine function. Consequently, the firing events of a representative neuron were plotted in a three-dimensional space.

Additionally, time points were captured at intervals of 10 msec for a duration of 8 sec (equivalent to one wheel rotation) starting from a specific point during the session. The phases of left touch, right touch and licking were calculated for each time point and converted using the cosine function. Finally, the movement trajectory for one wheel rotation was depicted in three-dimensional space by connecting those time points.

