## Supplemental Figures for "Rhythm Receptive Fields in Striatum of Mice Executing Complex Continuous Movement Sequences"

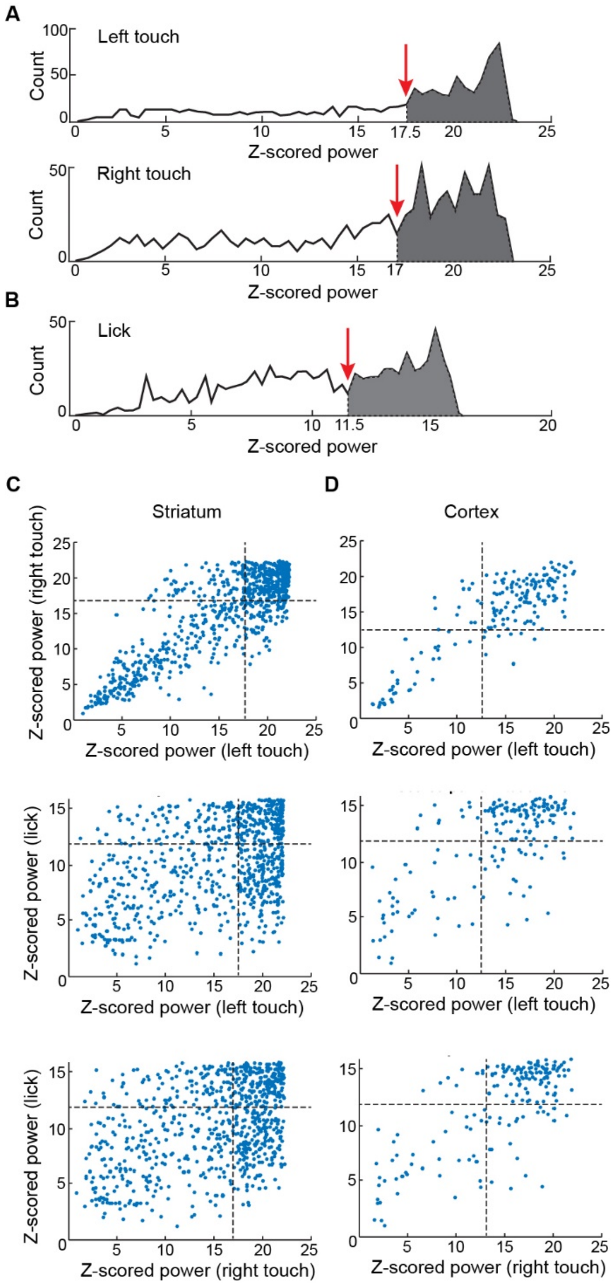


**Figure S1: Distributions of peak z-scored power, related to Fig. 2 and STAR Methods.**
A. To identify touch-responsive MSNs, we utilized the spectrogram obtained by performing a DFT on the cross-correlogram of the spikes aligned to touches. The spectrograms were z-scored, and the distribution of peak z-scored power of frequency band corresponding to left (top) and right (bottom) touches (2-3.5 Hz) are illustrated. The thresholds for each z-scored power are indicated by red arrows on the distributions of peak z-scored power, and MSNs exceeding the threshold are shown with gray shadings.
B. For lick-responsive MSNs, we determined the peak z-scored power of frequency band corresponding to licking (7-11 Hz) using the same method.

C and D. Z-scored power of individual striatal MSNs (C) and cortical neurons (D) corresponding to left and right touches (top), left touches and lick (middle), right touches and lick (bottom) are illustrated as scatter plots. The threshold for each z-scored power are indicated by dotted lines.

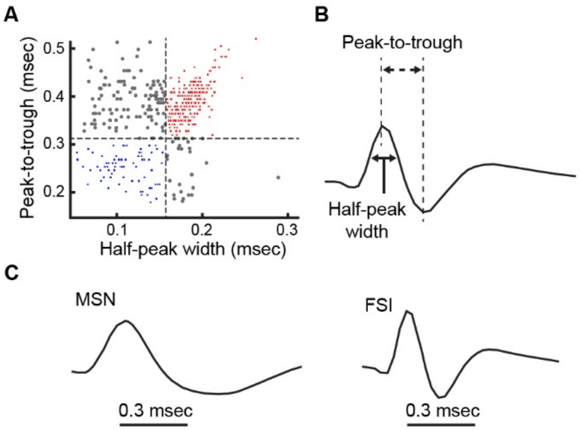


**Figure S2. Classification of each cell type, related to STAR Methods.**
A. Peak-to-trough and half-peak widths were calculated from the recorded waveforms, then units were classified as MSN (red; peak-to-trough width > 0.31 msec, half-peak-width > 0.16 msec), FSI (blue; peak-to-trough width < 0.31 msec, half-peak-width < 0.16 msec) and other cells (gray).
B. Illustration of peak-to-trough and half-peak widths in the waveform.
C. Representative waveforms of MSN and FSI.

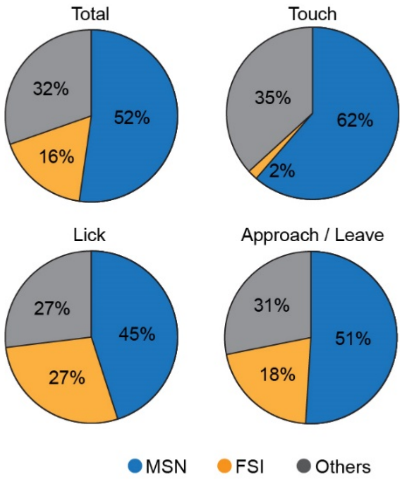


**Figure S3. The proportion of MSN, FSI and other neuron types, related to Fig. 2.**
The proportion of MSNs (blue), FSIs (yellow) and other neurons (gray) in all neurons recorded from all mice (Total), neurons that responded to touch (Touch), licking to the water spout (Licking), and approach to/leave from the water spout (Approach / Leave).

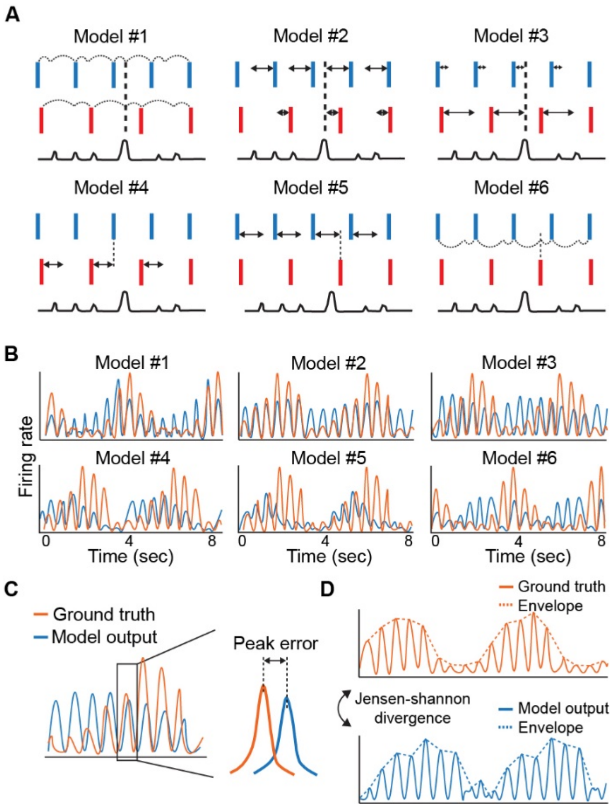


**Figure S4. Firing rate prediction using each hypothesis, related to Fig. 4.**
A. Conceptual diagram of six hypotheses.
**Hypothesis 1:** Striatal neurons have an optimal phase (relative time) to respond in each left and right cycle, and the firing rate changes according to the superposition.
**Hypothesis 2:** Striatal neurons have an optimal absolute spike-to-touch interval (time from firing to touch) to respond in each left and right cycle, and the firing rate changes according to the superposition.
**Hypothesis 3:** Striatal neurons have an optimal absolute touch-to-fire interval (time from touch to firing) to respond in each left and right cycle, and the firing rate changes according to the superposition.
**Hypothesis 4:** Striatal neurons have an optimal absolute right-to-left touch time (time from the touch of the right limb to the touch of the following left limb) to respond.
**Hypothesis 5:** Striatal neurons have an optimal absolute left-to-right touch time (time from the touch of the left limb to the touch of the following right limb) to respond.
**Hypothesis 6:** Striatal neurons have an optimal timing to respond when touch of the right limb occurred in a specific phase (relative time) within the left cycle.
B. Predicted (blue) and actual (orange) firing rates during running on the phase-changing peg-pattern based on each six model.
C. Evaluation index for prediction accuracy of each model (peak error) based on the distance of the peaks in the firing rate between the predicted (blue) and actual (orange) data.
D. Evaluation index for prediction accuracy of each model by JSD. Distance between the distributions obtained from envelopes of predicted and actual firing rates.

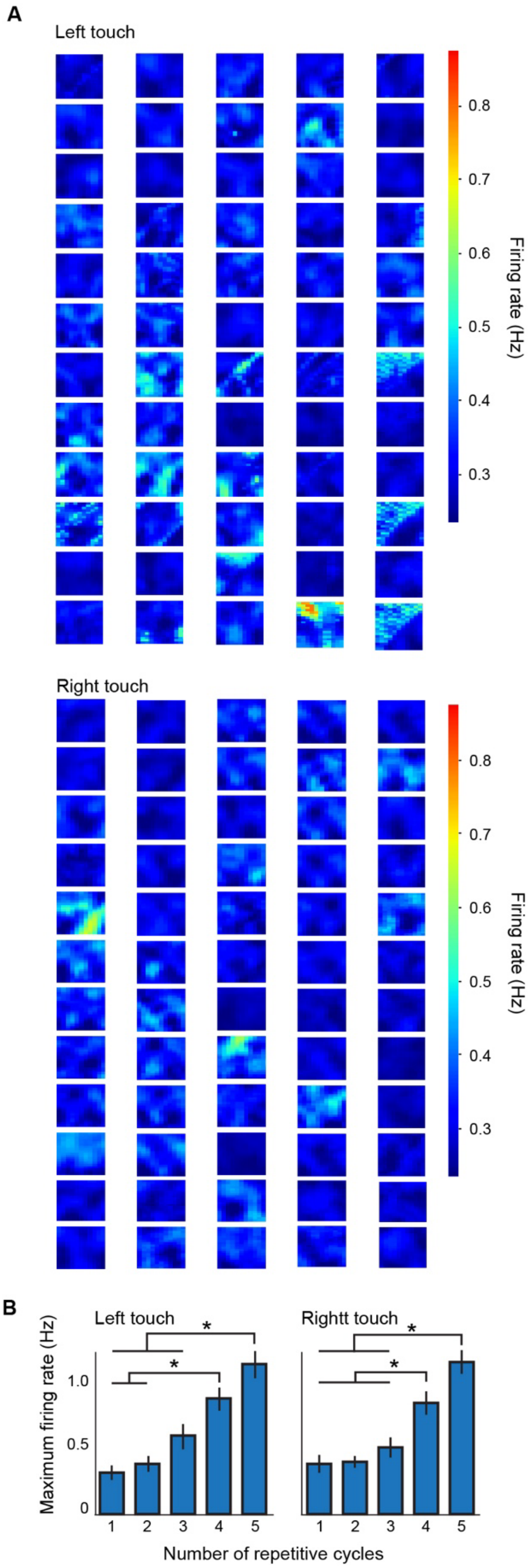


**Figure S5. Interval-phase firing characteristic map at low repetition state, related to Fig. 6.**

A. Heatmaps of interval-phase firing characteristics, calculated by window sets with various combinations of intervals and phases. The firing rate at the low repetition state (with 3 or less repetitive touch cycles) is shown in heatmap for each combination of interval and phase. The same neurons presented in Fig. 6B are shown.
B. The maximum firing rate of all touch-responsive MSNs, recorded from mice that ran on the phase-changing, interval-changing and complex peg-pattern, in relation to the number of repetitive touch cycles ($n = 112$ units, $p < 0.05$ by one-factor ANOVA; $^*p < 0.05$ by Bonferroni test).
